# Dynamic hippocampal and cortical signatures of naturalistic event memory

**DOI:** 10.64898/2026.09.06.749693

**Authors:** Marta Silva, Xiongbo Wu, Marc Sabio, Estefanía Conde-Blanco, Pedro Roldan, Antonio Donaire, Mar Carreño, Lluís Fuentemilla, Christopher Baldassano, Joshua Jacobs

**Affiliations:** Department of Neurology, University of Chicago, Chicago, USA; Department of Psychology, Ludwig-Maximilians-Universität München, Munich, Germany; Department of Cognition, Development and Education Psychology, University of Barcelona, Spain; Institute of Neuroscience (UBNeuro), University of Barcelona, Barcelona, Spain; Department of Neurology, Hospital Clínic de Barcelona, University of Barcelona, Barcelona, Spain; Fundació de Recerca Clínic Barcelona, Institut d’Investigacions Biomèdiques August Pi i Sunyer, Barcelona, Spain; European Reference Network EpiCARE, Barcelona, Spain; Department of Neurosurgery, Hospital Clínic de Barcelona, University of Barcelona, Barcelona, Spain; Bellvitge Institute for Biomedical Research, IDIBELL, Hospitalet de Llobregat, Spain; Department of Psychology, Columbia University, New York, USA

**Keywords:** Episodic memory, event segmentation, oscillations

## Abstract

Remembering temporally extended experiences requires integrating information over time into coherent event representations that can be stored and later retrieved. This process places changing demands on memory as an event unfolds: event representations must be reset at the beginning, connected into a unified model during the event, and stored in long-term memory at the end. We hypothesized that oscillatory dynamics, by coordinating communication across regions and organizing neural activity in time, provide a flexible mechanism for meeting these changing demands. However, most evidence comes from discrete-stimulus paradigms, leaving unclear how they support continuous experience. We recorded intracranial electroencephalography (iEEG) while participants watched a movie and later freely recalled it, and tested how power, phase synchrony and alignment at the beginning, middle, and end of each event related to subsequent memory. At beginnings, remembered events showed reduced gamma power and increased hippocampal–frontal synchrony. Within an event, memory was predicted by increasing temporal gamma power and hippocampal–temporal gamma synchrony, consistent with progressive binding of information. At event endings, memory was reflected by increased hippocampal theta power and phase resetting. Together these results provide a new perspective on how the brain supports memory in naturalistic settings, showing that it does not rest on one sustained encoding state. Instead, it requires temporal coordination of oscillatory states across brain regions and event phases, in which stable states support ongoing representation while transient, boundary-driven reorganization of hippocampal–cortical networks switches the system between storing the completed event and encoding the next.

**Significance Statement:** Theories about how brain rhythms support memory have assumed a single optimal brain state for encoding new information, based on studies presenting discrete items, such as words or images. Real-world experience, however, is continuous: events unfold over time and are encoded as connected memory traces. By recording electrical brain activity while patients watched and recalled a 50-minute film, we find that remembering an event depends not on one static encoding state but on a dynamic sequence of brain rhythms and hippocampal–cortical coordination that change as the event begins, unfolds, and ends. These findings show that previous discrete-item paradigms capture only a snapshot of one step in memory formation and reveal how the brain converts continuous experience into lasting memories.

## Introduction

Although episodic memory formation is often conceptualized and studied as a rapid process triggered by a discrete stimulus, real-world experience presents a fundamentally different challenge. Events unfold continuously, with information arriving moment by moment and overlapping across time and context. The brain must therefore determine which experiences should be bound together into a common memory representation and which should remain distinct. This requires a flexible mechanism for organizing continuous experience into temporally extended units that can later be retrieved as coherent episodes. We propose that the brain exhibits distinct neural signatures related to the segmentation of memory episodes, allowing us to extract distinctly remembered events from the continuous experience of our lives. This segmentation is reflected in how we remember: people recall past experience as a sequence of discrete events rather than as a continuous stream (1–3), and the structure of recall tracks the event structure of the original experience (4, 5). If memory is organized into discrete events, the brain must have neural processes that carve continuous experience into these memorable units. What are these processes, and how do they change as an event unfolds?

When a human experiences a new event, the brain initiates a new state that allows it to distinguish the newly begun event from the events that happened moments ago. This may involve a reset of working memory at the start of the event, which limits interference from the preceding event, and the use of schematic prior knowledge to rapidly build a template for the new event (2, 6). Next, during an event, incoming information must be processed and integrated into the evolving event model (7, 8). Finally, when an event boundary at the end of an event is detected, this is thought to be a critical moment for updating and encoding the current event model into memory (5, 9). Together, these stages reflect different functions of the memory system, which may be met by different neural processes, in different brain regions, at different moments. A central question is therefore how neural activity in different regions, and the coordination between them, supports memory encoding as these demands change. Because almost all prior work makes use of neuroimaging methods that are limited in temporal (e.g. fMRI) or spatial resolution (e.g. scalp EEG), our understanding of the specific neural mechanisms underlying these processes has been limited.

Direct brain recordings demonstrate some of the key neural mechanisms underlying memory processes. In particular, these studies showed that neuronal oscillations demonstrate many plausible mechanisms that can support the encoding of experiences at the different moments of an event. Neural oscillations, especially those in the theta and gamma frequency bands, may be informative about the processes that underlie event memory. These oscillations can simultaneously facilitate communication among distributed neural populations, provide a temporal framework for synaptic plasticity, by regulating how the phase of ongoing rhythms influences whether coincident activity potentiates or depresses synaptic connections (10, 11), and structure information processing over time (12–14). However, our current understanding of how neural oscillations support memory processing comes from discrete-stimulus paradigms, in which individual items (e.g., words, images, or objects) are presented one at a time, typically for a fixed duration and separated by relatively well-defined intervals. In these settings, a consistent spectral signature of successful memory encoding has emerged, consisting of increases in theta (15–18) and gamma power (19, 20), together with reduced alpha/beta activity (15, 17, 21–24), reflecting coordinated changes in local and network-level processing during the formation of lasting memories. This signature extends beyond local power: successful encoding of discrete stimuli is also accompanied by increased oscillatory synchrony, particularly in the theta (25–28) and gamma ranges (29, 30), supporting the view that memory encoding reflects coordinated interactions across distributed cortical and hippocampal networks. In discrete-stimulus paradigms, then, successful memory has largely been characterized by a single, stable oscillatory state.

How might the neural signature of memory generalize to temporally extended experiences? One possibility is that the neural signature found in discrete-stimulus paradigms carries over directly to continuous experience, so that recording an extended event into memory simply requires maintaining one favorable encoding state, marked by elevated theta and gamma power together with reduced alpha/beta (31). This account would predict that the overall prevalence or strength of this encoding state throughout an event dictate whether incoming information is incorporated into memory. Encoding-related activity should therefore not show systematic differences between the beginning, middle, and end of an event beyond those arising from changes in the information or task demands themselves. The complex structure of events in everyday life, however, suggests a second possibility: the demands placed on the memory system may change as an event unfolds. This means that encoding may not rely on a single, stable neural state, but rather on a sequence of states that dynamically regulate the brain’s readiness to process and retain incoming information.

These two accounts above make different, testable predictions. If oscillatory dynamics reveal the separate processes that organize continuous experience into discrete, memorable events, then neural oscillations should change systematically across different stages of an event, and these changes should predict whether a given memory is subsequently remembered. Testing this requires tracking oscillatory activity continuously across time and brain regions involved in event representations while participants engage with a naturalistic experience. Here we examined these issues for the first time with intracranial recordings using a naturalistic movie paradigm. A movie task gives us a unique opportunity to explore these dynamics, which a discrete stimulus cannot capture. We simultaneously recorded intracranial electrophysiological activity from the hippocampus, frontal cortex, and temporal cortex in patients undergoing treatment for pharmacologically intractable epilepsy while they watched the first 50 min of the first episode of BBC’s Sherlock. After the movie, participants completed a free verbal recall task, which allowed us to identify which events were later remembered and thus to probe how oscillatory dynamics during encoding relate to successful memory for each event. By comparing oscillatory power and synchrony between subsequently recalled and forgotten events at both beginning, middle and end of these events, we asked whether the temporal coordination and stability of neural activity across regions and event phases support the transformation of continuous experience into separable, retrievable episodes.

## Results

To investigate the oscillatory dynamics of naturalistic encoding, we recorded electrophysiological activity from intracranial electrodes implanted in seventeen epileptic patients while they watched the first 50 min of the first episode of BBC’s Sherlock (Fig. 1a). We focused on electrodes located in the hippocampus (58 channels), temporal cortex (422 channels) and frontal cortex (364 channels; Fig. 1b), regions with well-established involvement in episodic memory and event processing (3, 32–35). Subjects were then asked to freely recall the episode while their speech was recorded. The video stimulus was divided into 38 events, previously validated in (5), and for each participant we scored whether each scene was mentioned (remembered) or absent (forgotten) in their verbal recall. On average, we found that most participants showed good memory performance, successfully recalling 37% of the encoded events (SD = 4.02%). Moreover, they were accurate in maintaining the order in which the events were presented in the movie during recall (mean Kendall τ = 0.69, p < 0.001), similar to previous findings in healthy participants (5).

**Figure 1.**
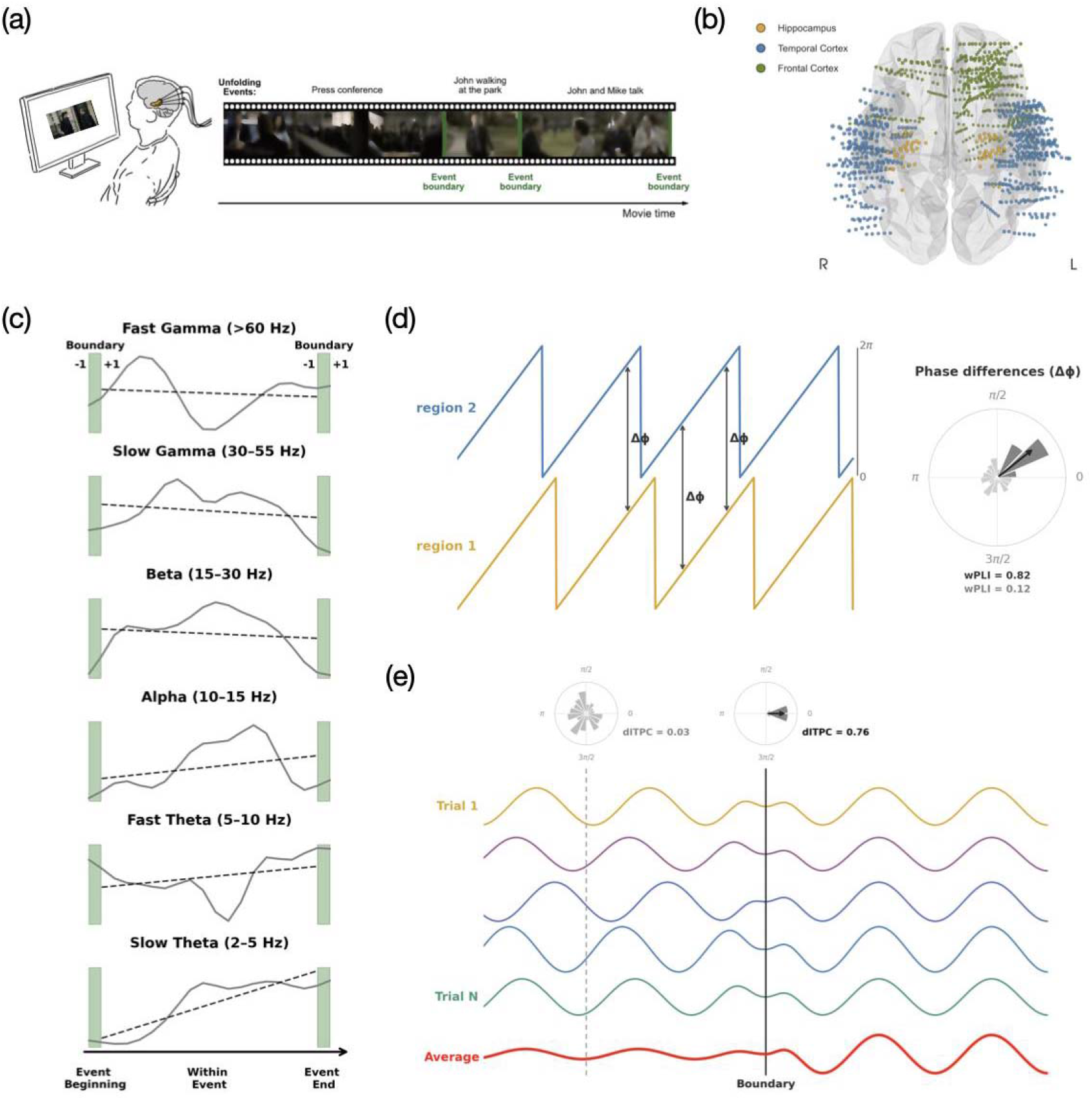
Experimental design, electrode placement and analysis schema. **a)** Experimental design. **b)** Hippocampal (yellow), temporal (blue) and frontal cortex (green) electrode localizations from all participants mapped into common space. **c)** Oscillatory power and synchrony were quantified within ±1s windows around event onset and offset (shaded green areas), and by the slope of activity across events (dashed black line), quantified by fitting a linear regression to the time-resolved values, for different frequency bands. **d)** Schematic of the phase synchrony analysis. For each channel pair, band-filtered signals from the two regions (region 1, yellow; region 2, blue) were used to extract the instantaneous phase difference (Δß) at each time point (arrows). The distribution of Δß across the analysis window is shown as a polar histogram, with the black vector indicating its resultant direction and magnitude. Phase differences that remain consistent and non-zero across the window yield high wPLI. **e)** Schematic of the phase reset analysis. Band-filtered traces from individual events (colored lines) are aligned to the annotated event boundary (black dashed line), with dots marking the instantaneous phase of each event at that time point. Polar histograms show the distribution of phases across events at a randomly selected time point (left, grey dashed line) and at the boundary (right). Dispersed phases yield low ITPC, whereas phase alignment across events at the boundary yields high ITPC (bottom). Observed values were evaluated against the null distribution obtained by shuffling boundary timings across events (grey shaded area).

### Oscillatory signatures of memory at the event beginning

Because the memory demands of an event change as it unfolds, we hypothesized that distinct oscillatory mechanisms support memory formation at its beginning, middle, and end. We designed distinct data analyses to separately probe these periods.

We first asked what neural state supports memory at the beginning of an event. In discrete-stimulus paradigms, the brain state immediately preceding an item predicts whether it will later be remembered. For example, medial temporal theta power before stimulus onset predicts encoding success (36, 37). This suggests that successful encoding depends on the system being in a receptive state when new information arrives. Similarly, at the onset of an event, the system must disengage from the preceding event and establish a new representation based on incoming external information. We therefore reasoned that a successful onset would reflect a preparatory state.

To test these hypotheses, we started by quantifying oscillatory power as the average of a ±1s window centered around event onset (Fig. 1c) and comparing subsequently recalled versus forgotten events using linear mixed-effects models (LMMs), fitted separately for each frequency band and brain region. We used LMMs because they allow us to dissociate the effects of memory for the subsequent event from those of the preceding event, while capturing variability arising from multiple sources, including participants, electrodes, and events. The β fitted from this LME model thus indicated the direction of any relations between the power of oscillatory signals and the success of memory encoding. Using this model thus revealed whether oscillations at specific frequency bands support the initiation of successful event encoding. In the hippocampus, recalled events were characterized by increased alpha power (β = 0.4099, p < 0.001) and reduced fast gamma power at event onset (β = - 0.1617, p < 0.001). In the temporal cortex, successful encoding was associated with increased power in the slower theta range (*β* = 0.2371, p < 0.001), but decreased beta power (β =-0.0708, p = 0.0528) and gamma power across both slow (*β* =-0.2498, p < 0.001) and fast gamma bands (*β* =-0.2534, p < 0.001). In contrast, in the frontal cortex memory was associated with broadly reduced power across most frequency bands (fast theta: *β* =-0.1780, p < 0.001; alpha: β =-0.2791, p < 0.001; beta: *β* = - 0.0912, p < 0.001; slow gamma: *β* =-0.080, p = 0.0197; fast gamma: *β* =-0.1774, p < 0.001), with the exception of slow theta power (Fig 2a). These power changes suggest that successful events begin with less local processing, raising the possibility that they might additionally depend on coordination between regions.

**Figure 2.**
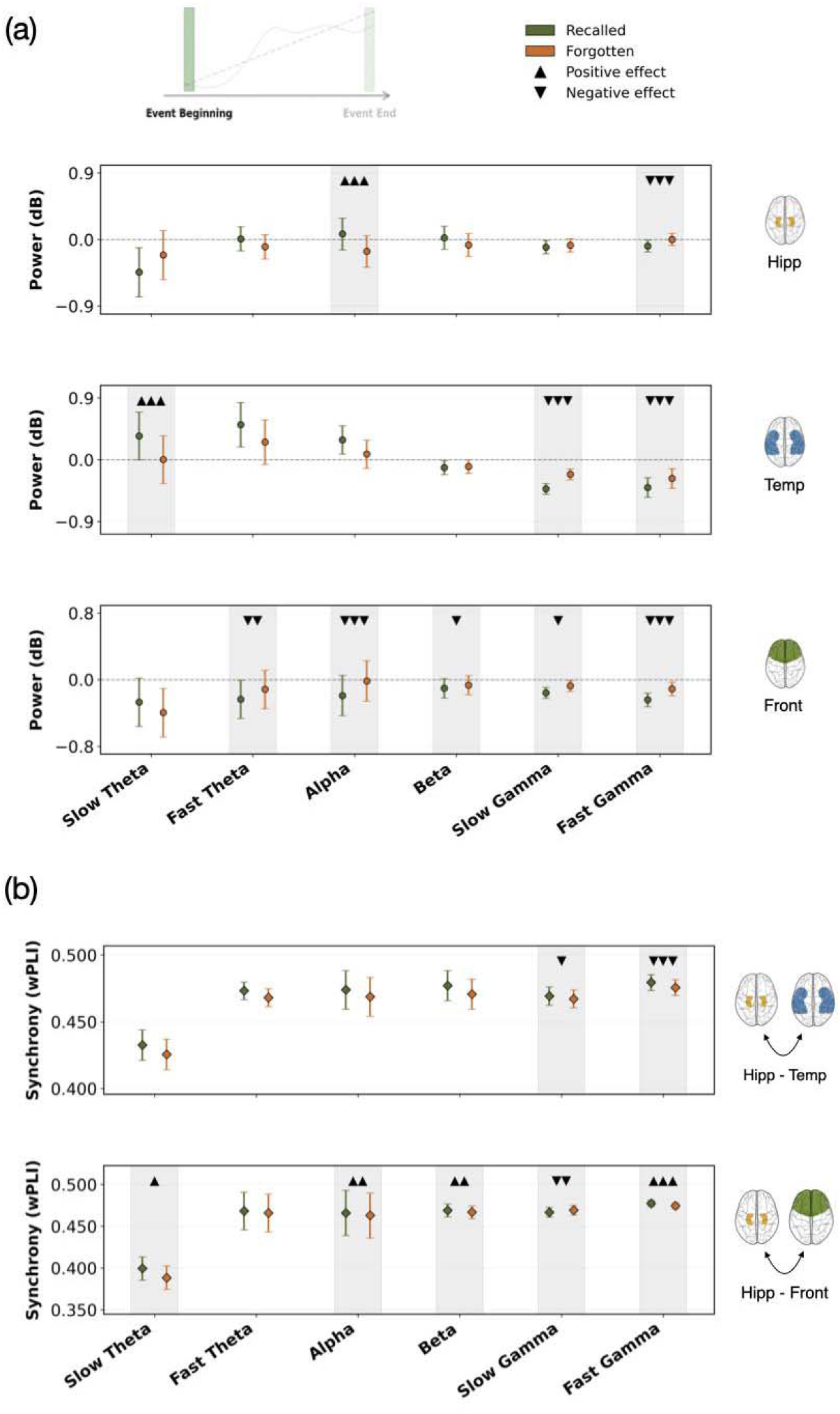
Power and synchrony memory effects at the beginning of events. Band-specific modulation of (**a**) power and **(b)** hippocampal–cortical synchrony as a function of memory for the upcoming event, with frequency bands along the x-axis. Green and orange markers show the estimated means for recalled and forgotten events, respectively. Estimated means were computed from the mixed-effects model as linear combinations of the fixed-effect coefficients: for each condition, the predicted value combines the intercept, the relevant memory effects and their interaction, with event duration evaluated at its mean. Error bars show 95% confidence intervals computed from participant-level data using the Cousineau–Morey method. Triangles mark significant memory effects: upward triangles indicate higher values for recalled than forgotten events, downward triangles indicate lower values, and one, two or three triangles denote p < 0.05, p < 0.01 and p < 0.001, respectively (Bonferroni-corrected across frequency bands and regions). Shaded grey areas highlight frequency bands with a significant effect. The inset schematic shows the analysis window (event onset).

Successful encoding of discrete items had also been shown to be reflected by increased hippocampal–cortical synchrony, particularly in the theta and gamma ranges (25, 26, 29). We therefore asked whether the receptive state at event onset extends beyond local power changes to coordination between the hippocampus and cortex. To test this, we next examined phase synchrony between the hippocampus and cortical regions, quantified as the weighted phase-lag index (wPLI) across all hippocampal–cortical channel pairs (Fig 1d). At event onset, hippocampus and temporal cortex showed increased beta-band coupling for subsequently recalled events (*β* = 0.0059, p < 0.001), accompanied by reduced synchrony in both slow (*β* =-0.0036, p = 0.0067) and fast gamma bands (*β* =-0.0037, p < 0.001). Hippocampus–frontal cortex interactions showed a different pattern, with subsequently recalled events exhibiting increased synchrony in most frequency bands (slow theta *β* = 0.0084, p = 0.0164; alpha β = 0.0095, p = 0.0014; beta *β* = 0.0063, p < 0.001; fast gamma *β* = 0.0023, p < 0.001), except for a decreased synchrony in slow gamma (*β* =-0.0038, p = 0.0029) relative to forgotten events (Fig 2b). Thus, onset-related coupling differed by cortical target and successful memory was therefore accompanied by strengthened low-frequency and beta coordination, particularly with the frontal cortex, rather than by uniformly increased gamma coupling.

Overall, these findings suggest that at the beginning of an event, successful memory is associated with specific spectral signatures, consistent with the establishment of an encoding-relevant neural state. The successful beginning of an event that is subsequently remembered is marked by long-range coordination across the brain, rather than increased local processing.

### Within-event oscillatory dynamics and subsequent memory

Having examined how events that are successfully remembered begin, we next asked what supports memory as the event unfolds, when the evolving neural representation must accommodate a continuous stream of sensory, semantic, and contextual information while preserving the coherence of what has already been encoded. The neural state supporting ongoing encoding must therefore balance the processing of newly arriving information with the maintenance and updating of the current event representation. Gamma oscillations have been consistently linked to local information processing and the active representation of information (Jensen, Kaiser and Lachaux, 2007; Fries, 2015), making them a plausible mechanism for the rapid processing and incorporation of continuously arriving information. Additionally, cortical activity during continuous experience has been shown to reflect the ongoing representation of events, with different regions maintaining representations over different temporal scales (Ranganath and Ritchey, 2012; Baldassano *et al.*, 2017; Yeshurun, Nguyen and Hasson, 2021). Gamma activity in the cortex could therefore reflect the ongoing processing and integration of event-specific information, while its coordination with the hippocampus may support the incorporation of these details into an evolving event representation. To test these hypotheses, next we examined how neural oscillations evolved during events.

A key challenge of naturalistic paradigms is that events vary substantially in duration, making direct temporal comparisons or aggregation across events difficult. To measure changes in neural signals during an event while accommodating their varying timecourses, we measured the linear trend of both power and phase synchrony of each oscillation within each individual event and extracted its slope (Fig 1c). Our findings indicate that successful and unsuccessful memory formation are associated with distinct ramping of oscillatory activity and hippocampal–cortical coordination during ongoing events. In the hippocampus, successful events exhibited decreases in slow theta (*β* =-0.3351, p = 0.0475) and beta (*β* = 0.1944, p = 0.0038) power relative to forgotten events. Similarly, in temporal cortex, recalled events showed more negative slopes in slower bands (slow theta *β* =-0.6288, p < 0.001; fast theta *β* =-0.4463, p < 0.001; alpha β =-0.3360, p < 0.001; beta *β* =-0.1006, p < 0.001) and stronger increases in the gamma band (slow gamma *β* = 0.2064, p < 0.001; fast gamma *β* = 0.2492, p < 0.001; Fig. 3a). Recalled events also exhibited greater increases in hippocampal–temporal fast gamma synchrony (*β* = 0.0025, p < 0.001) and steeper decreases in hippocampal–temporal alpha synchrony (β =-0.0044, p = 0.0113). In frontal cortex, forgotten events showed steeper increases in slow theta (*β* =-0.2614, p < 0.001) and beta (*β* =-0.0936, p = 0.0023) power, whereas recalled events showed stronger increases in alpha power (β = 0.2271, p < 0.001) and were associated with lower slopes of hippocampal–frontal slow theta (*β* =-0.0074, p < 0.001; Fig. 3b).

**Figure 3.**
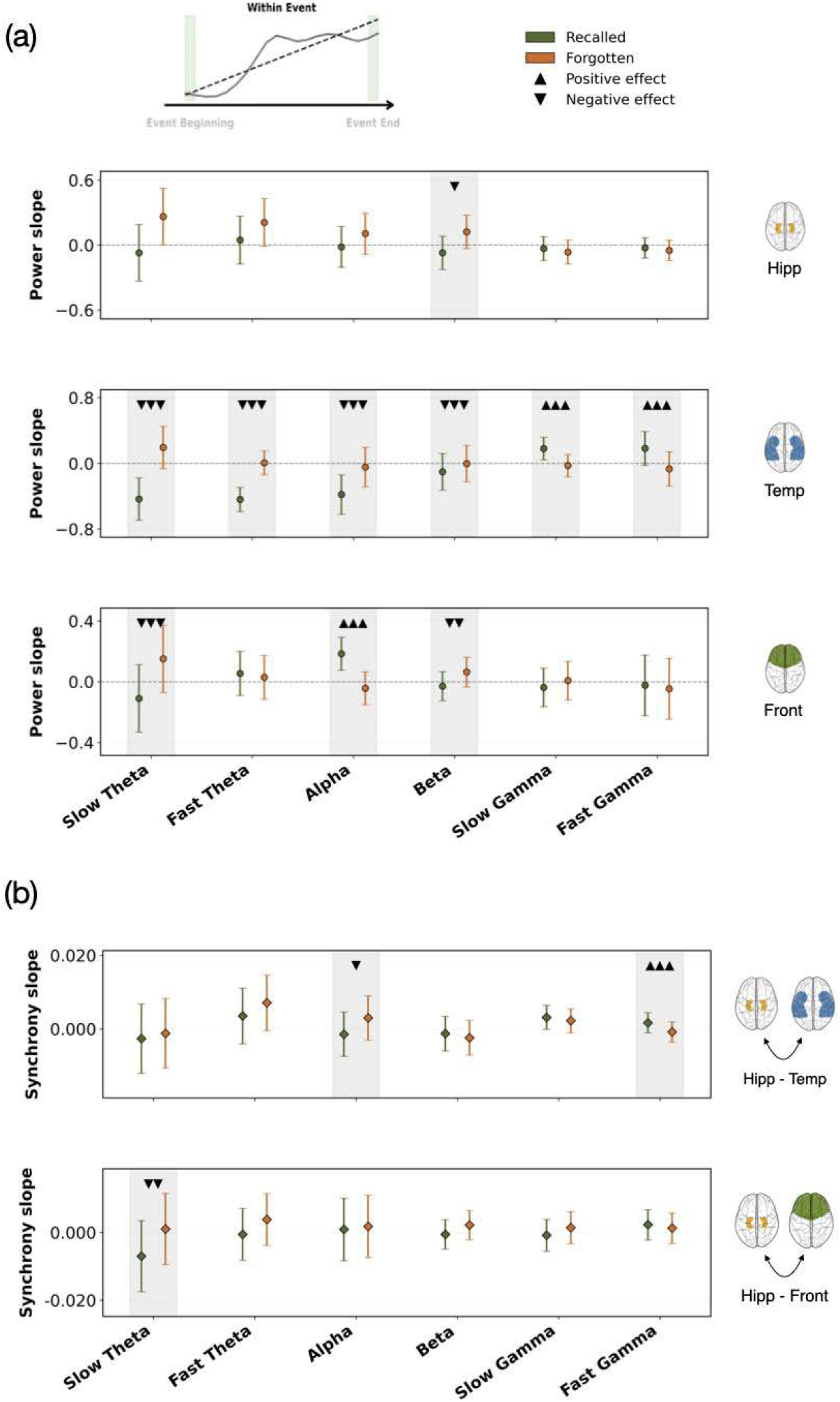
Within event slope effects on memory. Across events, band-specific modulation of (**a**) power and **(b)** hippocampal–cortical synchrony, quantified by fitting a linear regression to time-resolved values within each event, with the resulting slope used as an index of within-event change. Green and orange markers show the estimated mean slopes for recalled and forgotten events, respectively; positive slopes indicate an increase over the course of the event and negative slopes a decrease. Estimated means were computed from the mixed-effects model as linear combinations of the fixed-effect coefficients: for each condition, the predicted slope combines the intercept, the relevant memory effects and their interaction, with event duration evaluated at its mean. Error bars show 95% confidence intervals computed from participant-level data using the Cousineau–Morey method. Triangles mark significant memory effects: upward triangles indicate greater slopes for recalled than forgotten events, downward triangles indicate smaller slopes, and one, two or three triangles denote p < 0.05, p < 0.01 and p < 0.001, respectively (Bonferroni-corrected across frequency bands and regions). Shaded grey areas highlight frequency bands with a significant effect. The inset schematic shows the within-event slope analysed.

These findings indicate that successful memory formation was associated with distinct trends of oscillatory power and hippocampal–cortical synchrony during ongoing events. Where coordination occurs matters as much as whether it occurs, since rising hippocampal–temporal gamma synchrony accompanied successful recall of a particular event while rising hippocampal–frontal slow-frequency synchrony accompanied unsuccessful recall.

### Oscillatory signatures of memory at the end of an event

The end of an event is the last step in encoding the current event model into memory. This means the completed event must be integrated into long-term memory while a situation model for the next event is initiated, requiring a transient reconfiguration of communication between memory-related regions. These processes at event boundaries have been associated with increased hippocampal activity and rapid reinstatement of recently encoded information (3, 5, 34, 40). Accordingly, we have several predictions regarding how neural oscillations would behave in this period. Hippocampal theta is strongly implicated in episodic memory encoding and the organization of newly formed representations (16, 41). Hippocampal boundary-related increases in theta may therefore signal the termination and stabilization of the current event. Theta may speak to how the event is stored and shared across regions, but updating also requires that the ongoing event model itself be relinquished. Beta oscillations have frequently been associated with the maintenance of the current cognitive state and the propagation of top-down predictions (42, 43). Accordingly, decreases in frontal beta power at event boundaries may reflect the disruption of the existing event model and the need to revise ongoing predictions. To test these predictions, we therefore examined which oscillatory signatures were associated with successful encoding of the just-completed event by analyzing the averaged oscillatory power and phase synchrony of a ±1 s window surrounding event offset (Fig. 1c) and comparing recalled and forgotten events again by fitting linear mixed models separately for each frequency band and brain region.

In the hippocampus, recalled events were associated with increased theta-band power (slow theta *β* = 0.4715, p < 0.001; fast theta *β* = 0.3715, p = 0.0066) and reduced fast gamma power (*β* =-0.1267, p < 0.001). In contrast, the temporal cortex exhibited the opposite pattern, showing decreased theta power (slow theta *β* =-0.2443, p < 0.001; fast theta *β* =-0.3411, p <0.001), as well as decreased alpha power (β =-0.1884, p < 0.001), alongside increased slow gamma power (*β* = 0.0914, p < 0.001) for successfully recalled events. In the frontal cortex we see increased slow theta (*β* = 0.2166, p = 0.0037) alongside reduced beta (*β* =-0.1661, p < 0.001) and slow gamma activity (*β* =-0.0940, p = 0.0060; Fig 4a). Thus, at event offset, successful encoding was accompanied by opposite theta and gamma power profiles in the hippocampus and temporal cortex, together with reduced alpha power in the temporal cortex and reduced beta power in the frontal cortex. Considered jointly, these regions reproduce the spectral profile that had been previously observed to be associated with successful encoding of discrete items (31), but now additionally showing that the components of this signal were distributed across regions.

**Figure 4.**
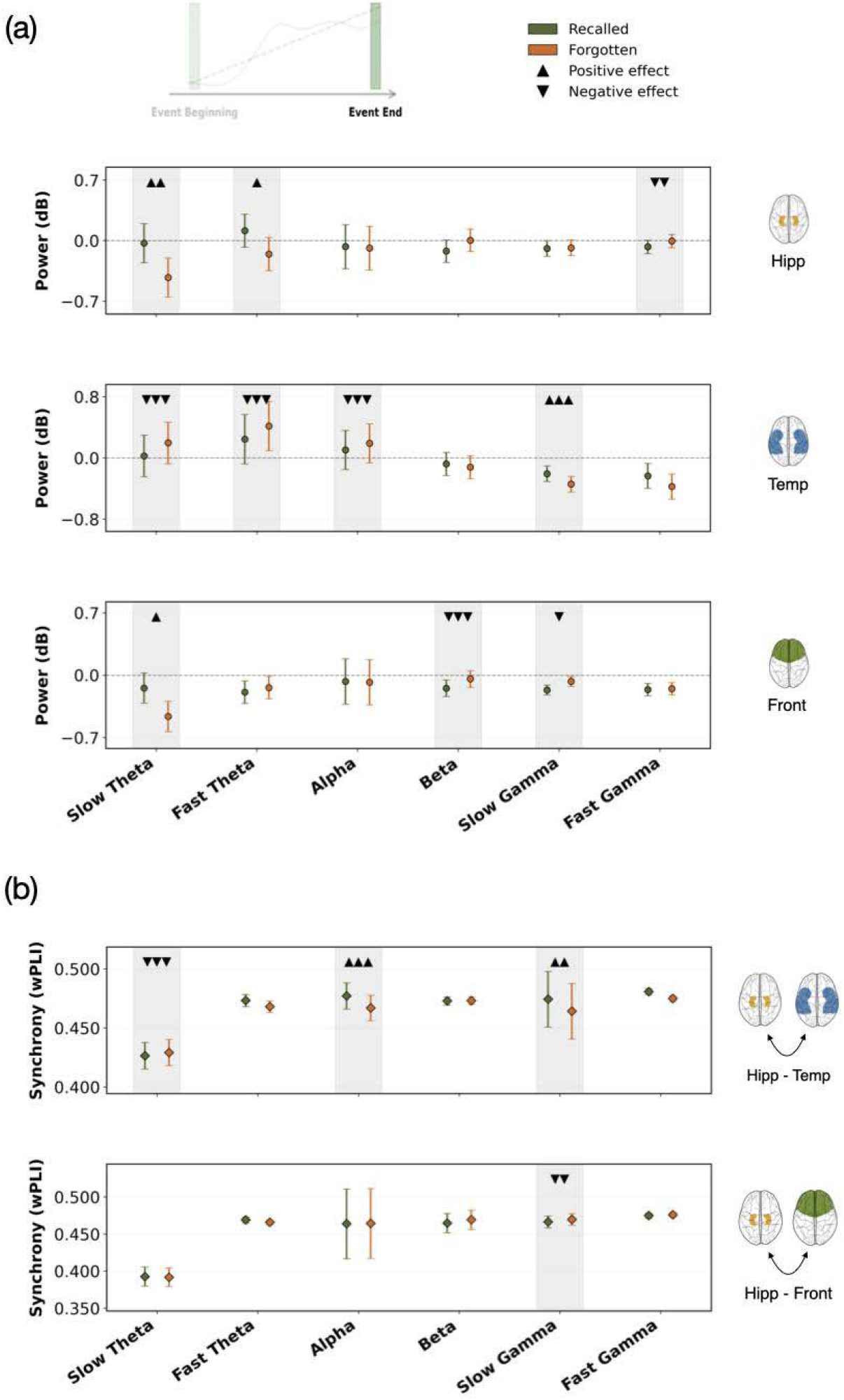
Power and synchrony memory effects at the end of events. Band-specific modulation of (**a**) power and **(b)** hippocampal–cortical synchrony as a function of memory for the just concluded event. Green and orange markers show the estimated means for recalled and forgotten events, respectively. Estimated means were computed from the mixed-effects model as linear combinations of the fixed-effect coefficients: for each condition, the predicted value combines the intercept, the relevant memory effects and their interaction, with event duration evaluated at its mean. Error bars show 95% confidence intervals computed from participant-level data using the Cousineau–Morey method. Triangles mark significant memory effects: upward triangles indicate higher values for recalled than forgotten events, downward triangles indicate lower values, and one, two or three triangles denote p < 0.05, p < 0.01 and p < 0.001, respectively (Bonferroni-corrected across frequency bands and regions). Shaded grey areas highlight frequency bands with a significant effect. The inset schematic shows the analysis window (event offset).

This pattern of results raised the question of whether these regional changes were also coordinated across regions. Communication between the hippocampus and cortical regions may be supported by increased gamma-band synchrony (29, 30). At event boundaries, enhanced hippocampal–cortical gamma coordination could support the transfer or integration of event-specific information as the preceding event is stored in long-term memory and the system transitions to a new context. To test this, we next examined phase synchrony between the hippocampus and cortical regions during event offset. We reasoned that if the hippocampus takes up a just-completed event from the cortical regions that hold its content, successful encoding should coincide with stronger gamma synchrony between the hippocampus and temporal cortex, where gamma power was elevated. We also expected altered hippocampal coupling with the frontal cortex, consistent with the release of the outgoing event model. Consistent with these predictions, synchrony between the hippocampus and temporal cortex was seen in slow gamma (*β* = 0.0046, p < 0.001) and alpha (β = 0.0105, p < 0.001) for recalled events, accompanied by reduced synchrony in slow theta (*β* =-0.0167, p < 0.001). Hippocampus–frontal cortex interactions showed successful encoding associated with reduced synchrony in slow gamma (*β* =-0.0046, p = 0.0011) frequencies relative to forgotten events (Fig. 4b). Thus, successful encoding of the just-completed event involved not only region-specific changes in power but also changes in hippocampal–cortical synchrony.

Taken together, these offset-related effects suggest that event boundaries constitute a brief but functionally important window for memory formation. Successful encoding appears to be supported by a coordinated shift in oscillatory dynamics across hippocampal and cortical regions, with increased theta activity in the hippocampus, increased gamma and reduced gamma in the temporal cortex and reduced beta activity in the frontal cortex.

### Differences in memory effects at event onsets versus offsets

Our central hypothesis was that successful event processing does not depend on a single neural state, but on oscillatory states that change as the phases of an event unfold. This predicts that the signatures of successful encoding at event onset should differ from those at event offset. Because several memory effects were present at both the beginning and the end of events, we next tested this directly, asking whether these boundary effects differed from each other, that is, whether they reflected position-specific processing of the event about to begin or the event just ending, rather than a general boundary effect. If boundary activity reflects a single general signal, memory effects should have the same direction at event onset and offset; if it reflects distinct cognitive demands, effects should differ by boundary position. We hypothesized that at least some signatures would be position specific. To test this, we tested a linear contrast comparing the two memory effects for each region and frequency band (Beg Vs End, Sup. Table 1). A significant contrast indicates that boundary activity predicts memory for the preceding and the following event differently.

For power, the contrast was significant for hippocampal slow theta (β = 0.6243, p = 0.0026) where increased power predicted memory for the preceding event but not for the following event. In the temporal cortex, memory effects differed between the preceding and the following event in most bands (slow theta β =-0.5129, p < 0.001; fast theta β =-0.4272, p < 0.001; alpha β =-0.2925, p < 0.001; fast gamma β = 0.2691, p < 0.001). Slow theta showed opposite effects for the two memory positions: higher slow theta power predicted memory for the following event, whereas the opposite pattern predicted memory for the preceding event. Fast theta and alpha power decreased specifically for the preceding event, and fast gamma power decreased specifically for the following event. In the frontal cortex, the contrast was significant for alpha (β = 0.1842, p = 0.0111) and fast gamma (β = 0.1195, p < 0.001), reflecting memory-related decreases specific to the following event.

Memory effects for hippocampal–cortical synchrony also differed for the beginning versus end of an event (Sup. Table 3). For hippocampal–temporal slow theta, reduced synchrony predicting memory for the preceding event but not the following event (β =-0.0096, p = 0.0178). Slow gamma synchrony showed opposite effects for onsets and offsets (β = 0.0082, p < 0.001): reduced synchrony predicted memory for the following event, whereas increased synchrony predicted memory for the preceding event. For hippocampal–temporal fast gamma, reduced synchrony was specific to the following event (β = 0.0023, p = 0.0239). For hippocampal–frontal interactions, memory-related increases in beta (β =-0.0068, p = 0.0150) and fast gamma synchrony (β =-0.0040, p < 0.001) were specific to the following event.

Together, these results indicate that during naturalistic behavior boundary-related activity does not reflect a single, general boundary signal. Some signatures, such as hippocampal slow theta power, temporal cortex fast theta and alpha power, and hippocampal–temporal slow theta synchrony, preferentially predicted memory for the event that had just ended. Others, such as temporal and frontal fast gamma power and hippocampal–frontal beta and fast gamma synchrony, were specific to the upcoming event. Temporal slow theta and slow gamma power and hippocampal–temporal slow gamma synchrony reversed in sign between the two events.

### Boundary phase resetting effects on memory

The preceding analyses show how much activity a boundary recruits and how strongly regions couple, but not when in the oscillatory cycle that activity falls. Oscillatory phase is known to impact synaptic plasticity: in the rodent hippocampus, stimulation at the peak of theta induces long-term potentiation, whereas stimulation at the trough induces depression (10, 44). A boundary that resets ongoing theta to the same phase across events could therefore align neural populations with the phase that favors encoding (45). Intracranial recordings in humans show that stimuli reset ongoing oscillations in neocortex and the medial temporal lobe during memory tasks (46, 47), and our newer research showed that phase alignment of hippocampal theta during encoding was causally relevant for memory (48, 49). We therefore hypothesized that boundaries would reset the phase of low-frequency oscillations, particularly in the hippocampus, and that stronger resetting would relate to memory, with the direction of the effect potentially depending on whether the reset served the event that had just ended or the one about to begin. Thus, we examined phase resetting at event boundaries in two steps.

We first tested whether event boundaries were accompanied by a reset of ongoing oscillatory activity and then asked whether the magnitude of this reset predicted memory for the event that was beginning or for the event that had just ended. To quantify this, we computed phase consistency across event boundaries for each channel (inter-trial phase coherence, ITPC) for frequencies below 10Hz and compared it against a surrogate null distribution (Fig 5a). We observed significant phase resetting in both the hippocampus (p = 0.0012; centered around ∼2 Hz) and temporal cortex (p < 0.001; centered around ∼3 Hz), whereas no significant reset was observed in the frontal cortex (Fig 5a). The resetting frequencies (∼2–3 Hz) fall within the slow theta range most consistently linked to episodic memory formation in human intracranial recordings (16), placing the effect in the band where a realignment of timing would be expected to be mnemonically consequential.

**Figure 5.**
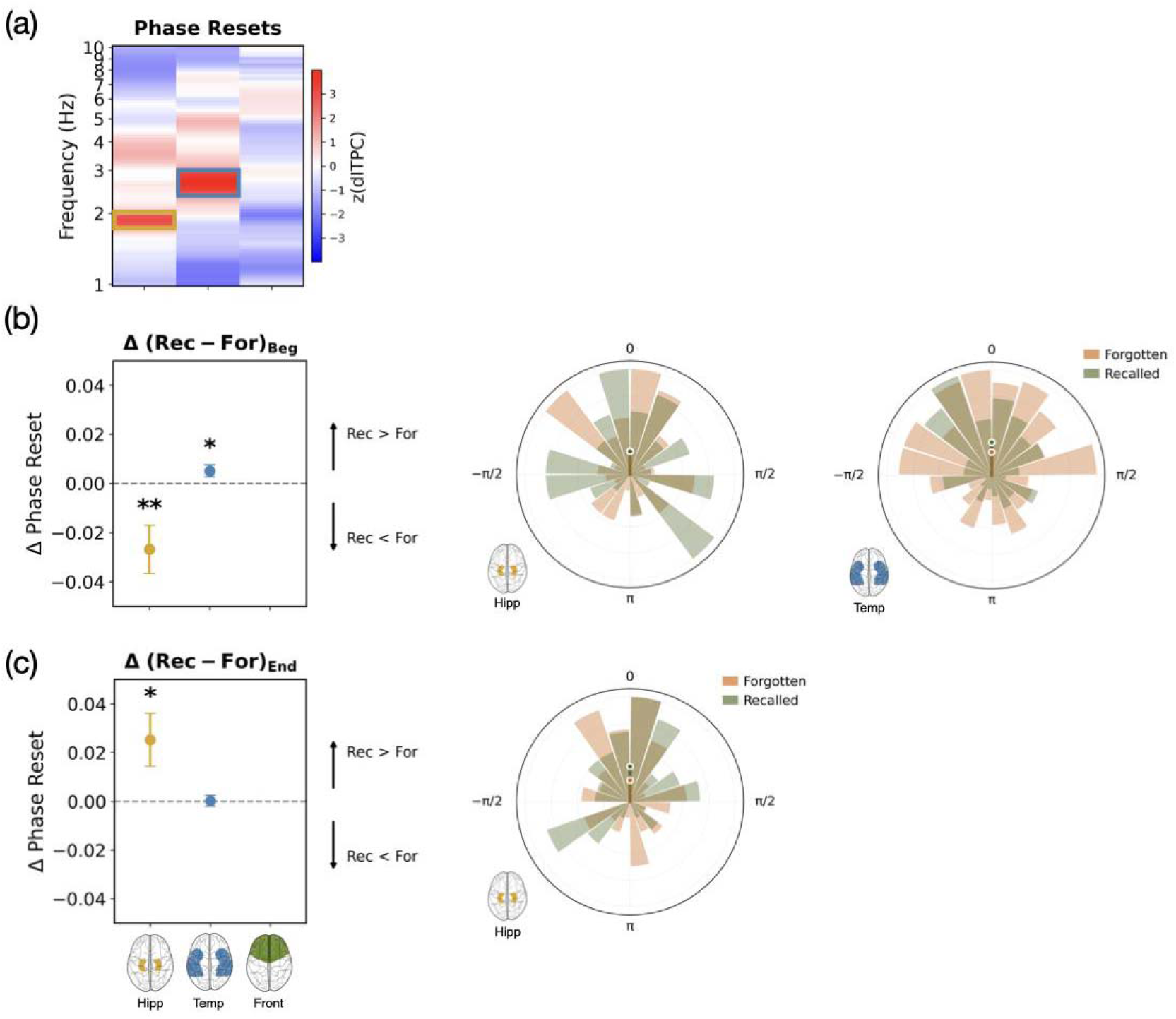
Phase reset memory effects. **(a)** The consistency of theta phase resets at event boundaries was measured with debiased ITPC and compared to a permutation-based null distribution. Clusters showing significant resetting to a consistent phase were identified in hippocampus (around 2Hz) and temporal cortex (around 3Hz). **(b)** Computing ITPC at these frequencies separately for the beginnings of events that were later recalled or forgotten shows significantly greater resetting for forgotten events in the hippocampus and remembered events in the temporal cortex. The polar plots show the aggregated angles at the boundaries across all participants for recalled (green) and forgotten events (orange) aligned to their preferred phases for the regions with significant phase resets. **(c)** Computing ITPC for the endings of events that were later recalled or forgotten shows significantly greater resetting for remembered events in the hippocampus. The polar plot shows the aggregated angles at the boundaries across all participants for hippocampal recalled (green) and forgotten events (orange) aligned to their preferred phases. ITPC values are scaled and plotted in the polar plots for visualization purposes, separately for recalled (green) and forgotten (orange) events.

**Figure 6.**
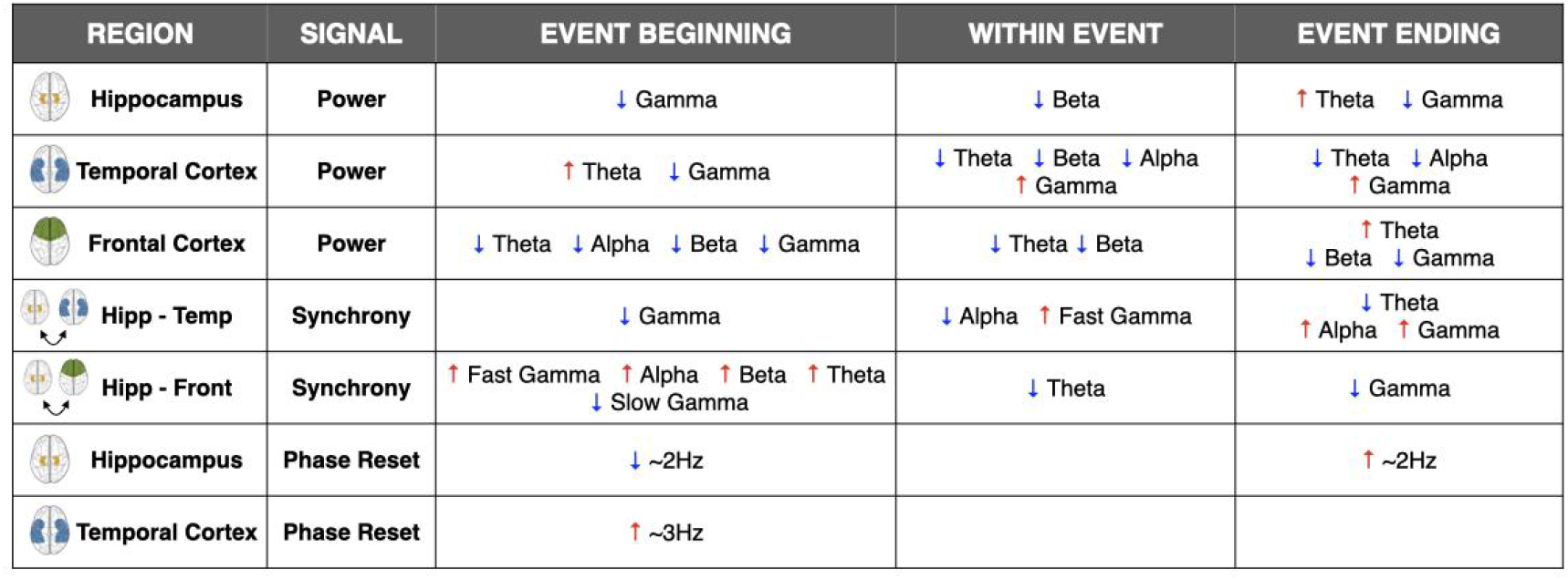
Summary of the memory effects divided by type of analysis, regions and event moment.

We then related the magnitude of boundary-related phase resetting to memory for the upcoming event. In the temporal cortex, stronger phase resetting predicted memory for the upcoming event (paired t-test, t_422_ = 2.0472, p = 0.0412; Fig 5b), whereas in the hippocampus it was associated with reduced memory for the upcoming event (t_57_ =-2.7347, p = 0.0083; Fig 5b). The hippocampal and temporal cortical effects were opposite in sign, indicating that a boundary does not elicit a single network-wide reset but region-specific realignments with opposing consequences for what is remembered.

Then, we investigated whether phase resetting at event boundaries contributed to memory for the just-concluded event. Here, we observed a memory-related effect specific to the hippocampus, where stronger phase resetting at event offset was associated with successful recall (t_57_ = 2.3122, p = 0.0244; Fig 5c) of the just concluded event. This positive effect of resetting for event offsets was significantly different from the negative effect at event onsets (t_57_ = 3.7219, p < 0.001), confirming that hippocampal phase resetting relates to memory in opposite directions depending on whether it supports the event that is beginning or the one that has just ended. These findings suggest that boundary-related hippocampal phase reorganization play a key role in stabilizing and storing recently encoded event representations.

The phase-reset findings show that this boundary-related reorganization also involves the timing of ongoing oscillations: beyond changes in oscillatory amplitude and inter-regional coupling, successful memory was related to how strongly ongoing activity was realigned at the boundary, and this relationship depended on both the region and the event being remembered. In the hippocampus, stronger resetting predicted better memory for the just-concluded event but poorer memory for the upcoming one, whereas in the temporal cortex stronger resetting predicted memory for the upcoming event. This suggests that the end of an event may trigger a transient reorganization of hippocampal timing that favors stabilizing the just-formed episodic representation at the expense of encoding the next one, while temporal cortical resetting may help initiate the representation of the new event.

## Discussion

In this study we set out to investigate the oscillatory dynamics of event encoding and how they might support both the integration of information during the encoding of an event. Because most of what is known about oscillatory contributions to episodic encoding comes from paradigms in which a discrete stimulus is presented and subsequently tested, a central question is whether the signatures established in those paradigms scale to experiences that unfold over tens of seconds. Overall, our results indicate that successful encoding was associated with distinct, frequency-specific oscillatory signatures across hippocampal, temporal, and frontal regions, as well as differentiated patterns of inter-regional synchrony throughout the event. In particular, the data suggest that oscillatory signatures supporting the encoding of a new event differ from those supporting the storage or stabilization of the just concluded event. The onsets of successfully encoded events showed reduced gamma activity in all regions, increased temporal theta, and reduced frontal activity. Additionally, the hippocampus showed increased beta synchronization with the temporal cortex and extended synchronization with the frontal cortex within most frequency bands. Within an event, distinct trajectories of oscillatory activity and hippocampal–cortical synchrony were associated with successful encoding, such as increased temporal gamma activity and hippocampal–temporal synchrony. Finally, at the end of an event, increased theta activity, phase resets, and reduced gamma activity in the hippocampus supported memory for the just-concluded event. Temporal cortical activity showed an opposite pattern: gamma power and synchrony with hippocampus was higher, while theta power and synchrony were reduced. In the frontal cortex, lower beta power at event boundaries was associated with better memory. Together, these results suggest that episodic memory formation during naturalistic experience depends on a dynamic division of labor across brain regions and frequencies, in which stable oscillatory states support ongoing representation, and transient, boundary-driven reorganization of hippocampal–cortical networks enable the switch between consolidation of past events and encoding of new ones.

Event boundaries provide an opportunity to enter a preparatory state to start encoding new information and establish a new event representation in working memory (2, 3, 34). Our results indicate that the initial stage of an event is characterized by coordinated neural dynamics that prepare distributed hippocampal-cortical networks for encoding upcoming information. We find generalized decreases in gamma activity, an increase in temporal theta power and an almost broadband decrease in frontal activity for events that were later recalled. This suggests a transient downscaling of ongoing local processing, potentially reflecting a reset of ongoing representations to accommodate new incoming information. In contrast to the increased high-frequency activity that commonly accompanies successful encoding of discrete items (19, 50), successful memory at event onset was associated with reduced local activity. This suggests that the onset of an extended event engages a different encoding regime from that observed for isolated stimuli, in which memory formation may depend less on sustained local processing and more on the establishment of a new event context. Despite this local suppression, network-level coordination increases, with almost broadband synchrony between frontal cortex and hippocampus predicting better subsequent memory. This suggests that transient increases in long-range coordination contribute to establishing new event representations or resetting a working memory model across event transitions (51). Together, these findings indicate that successful encoding of upcoming experiences relies on coordinated interactions between cortical regions and the hippocampus at the moment a new event begins.

Once an event model has been established, successful encoding requires its continued maintenance and updating as information unfolds over time (3, 52). Rather than repeatedly resetting network activity, oscillatory dynamics during ongoing experience are likely to support the continuous integration of sensory, semantic, and contextual information into a stable event representation. Previous work suggests that temporal cortex represents perceptual and conceptual aspects of ongoing events (32), whereas the hippocampus binds these features into episodic representations (3, 5, 34, 40). Within an event successful memory was associated with increasing gamma-band activity and synchrony in the temporal cortex, consistent with a possible role for gamma oscillations in supporting the representation and integration of event-specific information (14, 38). The synchrony findings further suggest that successful encoding depends on selective patterns of hippocampal–cortical communication. Recalled events showed stronger increases in hippocampal–temporal fast gamma synchrony, which may reflect enhanced interactions between the hippocampus and temporal cortex that support the binding and integration of event features into coherent memory representations (53). In contrast, forgotten events were characterized by progressively increasing beta power across hippocampal, temporal, and frontal regions, together with stronger trajectories of hippocampal–frontal synchrony in slow theta, potentially reflecting a non-optimal encoding state with continued updating, contextual integration, attentional fluctuations, or other processes that compete with the formation of durable event representations. This interpretation is consistent with recent work showing that hippocampal–neocortical interactions during the middle of events are detrimental to subsequent memory performance, both in computational models (54) and in empirical fMRI evidence (55). A limitation of our approach is that the temporal changes in oscillatory activity within events were summarized using only a linear regression slope. Neural activity during naturalistic events is likely not evolving in a linear fashion: event representations may be characterized by transient fluctuations, nonlinear trends or temporally localized bursts of oscillatory activity that cannot accurately be captured by a single linear parameter. Future work could employ more sophisticated approaches such as nonlinear modeling to better characterize within event temporal dynamics.

At the end of an event, the current event model must be finalized and recorded before the next event begins, preserving the information accumulated during the event while preventing interference between events (2, 56, 57). Event segmentation theories propose that this transition marks the moment at which one event representation is finalized and the just-encoded information stored into long-term memory (3, 5, 34, 40). Our hippocampal results support a particularly important role of theta dynamics in mediating this process. Overall, successful memory was associated with higher theta activity at boundaries together with decreasing activity throughout the event, suggesting that hippocampal processing relevant for memory is concentrated around moments of event transition rather than sustained continuously during ongoing experience. More specifically, theta activity appeared especially beneficial for memory of the just-concluded event, consistent with evidence that hippocampal theta supports successful episodic encoding and the organization of newly formed representations (16, 41). In the context of event segmentation, these dynamics may further reflect post-boundary stabilization processes that support the consolidation of the preceding event into a coherent memory trace (3, 34, 56). An additional open question pertains to the functional distinction between slow and fast theta oscillations. Previous work has proposed partially dissociable roles for these frequency ranges (58), with slow theta often linked to episodic memory processing (16, 59), and fast theta more strongly associated with online processing, including working memory, novelty detection, and sensorimotor engagement (60, 61). Although both theta bands contributed to successful memory formation at event boundaries in the hippocampus, directly comparing memory for the preceding and the following event suggests that they did so in different ways. Slow theta power was event-specific: it predicted memory for the just-concluded event but not for the upcoming one, as would be expected of a process that stabilizes the content of a particular episode. Fast theta power, in contrast, did not significantly differ from the effect predicting the upcoming event, consistent with a more general boundary-related state, such as heightened attention, novelty detection or updating, that benefits encoding regardless of which event is being stored. This dissociation aligns with the proposed link between slow theta and episodic memory processing, and with the association of fast theta with online processing. Future work will be required to determine whether these frequency bands reflect fully distinct mechanisms or complementary components of a shared process supporting the encoding and stabilization of continuous experience.

The phase-reset findings further reinforce this and add a dimension that power and phase synchrony alone cannot provide: beyond how much activity a boundary recruits, they show that a boundary reorganizes the precise timing of hippocampal activity. Hippocampal slow theta phase resets at event boundaries predicted better memory for the preceding event but poorer memory for the upcoming one, suggesting that boundary-triggered theta reorganization may preferentially facilitate the storage of recently encoded information rather than preparation for new encoding. This asymmetry provides an oscillatory counterpart to the offset-locked hippocampal responses that predict memory for the just-completed episode in fMRI (3, 34): the end of an event appears to be registered in the hippocampus not only as an increase in activity, but as a realignment of its intrinsic timing. These results further converge with recent single-neuron evidence that theta phase precession emerges following cognitive boundaries during movie watching and carries information about encoding success beyond firing rates (62). Such phase resetting may reflect a mechanism through which the hippocampus temporally aligns neural populations, facilitating hippocampal–cortical communication at event boundaries and promoting the stabilization of completed event representations in memory (47, 63).

In contrast to the heightened theta activity, successful memory was also associated with reduced gamma activity at event boundaries in the hippocampus, particularly reduced fast gamma power. This finding is especially notable given the concurrent observation of increased ripple activity (64) in a subset of this dataset, highlighting a potential functional distinction between hippocampal ripples and broadband gamma activity. Whereas ripples are thought to reflect highly coordinated replay and consolidation-related processes (65–68), elevated broadband fast gamma may instead index local excitatory processing or interference that is less beneficial for stabilizing recently encoded events. Together, these findings suggest that hippocampal event-boundary processing may rely on a transition toward slower, temporally coordinated theta states and ripple-related mechanisms, rather than increased broadband high-frequency activation.

Critically, the hippocampal boundary signature contrasts with the dynamics observed in the temporal cortex. This region showed the opposite pattern of activity, with increased gamma and reduced theta power. This regional dissociation suggests a division of labor across the hippocampal-temporal network. Considered jointly though, the two regions reproduce the spectral profile associated with successful encoding of discrete items (i.e theta and gamma increases alongside reduced alpha/beta power, Broitman and Kahana, 2026) which is typically observed within a single region. Continuous experience may therefore preserve the canonical encoding signature while distributing its components across the network and concentrate it at the moments when an event is completed. Whereas hippocampal theta and phase resetting may signal the termination of the current event and coordinate its stabilization, gamma activity in temporal cortex may sustain the perceptual and semantic representations that constitute the content of the event being consolidated (32, 52, 69–72). Rather than reflecting competing mechanisms, these oscillatory signatures may represent complementary computations operating in parallel: hippocampal theta organizing when an event should be segmented, and temporal gamma representing what information is preserved across that transition. Additionally, the effects of theta and gamma across regions raise the possibility of a phase–amplitude coupling (PAC) mechanism operating at event boundaries. Gamma bursts tend to occur at preferred phases of the theta cycle, allowing neuronal ensembles to be activated within discrete temporal windows (73, 74). This temporal organization has been proposed to facilitate communication between brain regions, coordinate information processing across multiple timescales, and support the sequential representation of information (51, 75, 76). At event boundaries, hippocampal theta might provide a temporal scaffold that organizes cortical gamma activity (53), but with distinct functional consequences depending on whether the system is stabilizing past information or transitioning toward new encoding demands.

Our frontal cortex results also resonate with previous literature on cortical dynamics during memory formation, which has consistently linked successful encoding to reductions in alpha/beta power in prefrontal regions (15, 21–24). These decreases are typically interpreted as reflecting a release from functional inhibition and a shift toward more efficient representational processing within cortical systems (21, 77). In our data, frontal alpha and beta were both reduced at event onset, and frontal beta was reduced at event offset, in line with this discrete-item signature. At onset, reduced frontal alpha may additionally reflect greater engagement in attentional control, monitoring, or organization of the incoming event. Lower frontal beta power at event boundaries was also associated with better memory for the upcoming event. This suggests that beta desynchronization in the frontal cortex may reflect a more general release from top-down maintenance of the current cognitive set (42) at boundaries, supporting a global reconfiguration state that benefits both encoding of past information and preparation for subsequent encoding. Boundaries are thought to arise when predictions about ongoing experience fail, triggering updating of the current situation model and reorganization of neural representations (2, 3). Accordingly, decreases in frontal beta at event boundaries may reflect the disengagement of the previous event model, allowing the system to update predictions and transition into a new event representation

Together, our findings support a model in which event boundaries act as critical transition points that reorganize hippocampal–cortical network dynamics to balance competing mnemonic demands. Importantly, the results also suggest that successful episodic memory depends not simply on stronger oscillatory activity, but on the appropriate temporal coordination and stability of oscillatory states across brain regions and event phases. Rather than reflecting a single encoding state that persists throughout an experience, as might be expected from previous research using discrete-item paradigms, successful memory in continuous experience is characterized by distinct oscillatory dynamics at event onset, within an event, and at its offset. Together, they point toward a dynamic interplay in which hippocampal synchronization mechanisms support episodic binding and encoding, while cortical desynchronization supports flexible information representation, with event boundaries serving as privileged moments for shifting between these complementary memory processes. Although the observed effects suggest an important role for hippocampal-cortical interactions, it remains to be explored how the current mechanisms extend to other regions involved in event encoding. Future studies with broader electrode coverage should investigate the contribution of additional areas to provide a better comprehensive account of the large-scale oscillatory dynamic supporting memory formation during continuous experiences.

## Materials and Methods

### Data Collection

We tested 17 human subjects (8 females, age range 18–60 years, mean ∼34.5 years), an extension of the Silva *et al.*, 2025 dataset, who were undergoing treatment for pharmacologically intractable epilepsy at Hospital Clínic—IDIBAPS in Barcelona. Prior to performing the task, all participants were thoroughly briefed on the specificities of the task, ensuring they had a comprehensive understanding of the objectives, procedures, and potential risks involved. Each participant was provided with a consent form, which they attentively reviewed and signed, demonstrating their informed consent to participate in the study. The study was approved by the hospital Ethics Committee. Patients were surgically implanted with intracranial depth electrodes for diagnostic purposes to isolate their epileptic seizure focus for potential subsequent surgical resection. Antiseizure medications were reduced to record seizures during the patient’s hospital stay. Cognitive testing was performed ≥7 h after the last seizure. The exact electrode number and locations varied across subjects and were determined solely by clinical needs. The recordings were performed using a clinical EEG system (Natus Quantum LTM Amplifier) with a 2048 Hz sampling rate and an online band-pass filter from 0.1 Hz to 4000 Hz. Intracerebral electrodes (Microdeep, DIXI Medical) were used for recordings. Each multielectrode had 8 to 18 contacts, spaced 5 mm and 1 to 2 mm long with a diameter of 0.8 mm.

### Experimental Design

The experiment was conducted in a sound-attenuated room in the hospital, with participants sitting upright in a comfortable chair or on their bed. Participants were asked to watch the first 50 min of the first episode of BBC’s Sherlock, dubbed in Spanish, as done previously (64). Participants were informed that a subsequent recall memory test would follow. After the movie, some time was given to rest (5–10 min) before the test began. During the test, they were asked to freely recall the episode without cues while being recorded using an audio recorder placed on the overbed table next to the laptop computer. The audio files were later analyzed to access the participants’ length of the recall. The experimental design was implemented using PsychoPy (78) and presented on a 13-inch portable computer, placed on the overbed table at approximately 60 cm distance in front of the patients.

### Event Boundary Annotations

The event model validated in previous research from the authors (Silva et al. 2019) was used for the current analysis. This model is composed by 38 events (minimum = 4 s, maximum = 444 s, and mean = 76.02 s) and it was constructed by having six external participants annotate the temporal point at which they felt “a new scene is starting; these are points in the movie when there is a major change in topic, location or time”. The final model was built based on boundary time points that were consistent across observers. To find a statistical threshold of how many observers should coincide in a given time point to be different from chance in our data, we shuffled the number of observations 1000 times and created a null distribution of the resulting coincident time points. An α = 0.05 as a cutoff for significance indicated that boundary time points at which at least 3 observers coincided in (considering 3 s as a possible window of coincidence as in Baldassano et al. 2017) could not be explained by chance.

### Verbal Recall Analysis

The audio files from the free verbal recall were analyzed by a laboratory member who was a proficient Spanish speaker, using the list of events from the event model mentioned in the previous section. An event was counted as recalled if the participant described any part of that scene. To statistically assess whether the order of events during movie watching was preserved during free recall, we computed Kendall rank correlation coefficients between each individual event temporal order and a simulated correct linear order. A positive Kendall tau coefficient close to 1 indicates that the encoded temporal order of the events was highly preserved during their recall.

### Electrode Localization and Selection

The presence of electrodes in the respective brain areas was assessed with the examination of a computed tomography (CT) and preoperative Magnetic Resonance Imaging (MRI) T1 scans. Electrode localization was performed using the YAEL (Your Advanced Electrode Localizer) pipeline implemented in the RAVE toolbox (79, 80). Pre-implantation T1-weighted MRI scans and post-implantation CT images were co-registered, and electrode contacts were manually identified using the interactive 3D visualization interface. Electrode coordinates were projected onto each participant’s cortical surface reconstructed with FreeSurfer, and anatomical labels were assigned based on standard atlas parcellations. Selection of channels was done in native space to prevent errors due to distortions.

For the present analyses, we focused on electrodes located in the hippocampus (58 channels), temporal cortex (422 channels) and frontal cortex (364 channels; Fig. 1b). These regions were selected a priori due to their well-established involvement in episodic memory and event processing. The hippocampus is critical for forming and updating event representations and binding information across time (3, 34), while the temporal cortex supports the representation of semantic and perceptual content relevant to events (3, 32). The frontal cortex contributes to higher-order control processes, including prediction, organization, and the evaluation of event structure (33, 35, 81). Together, these regions form a core network supporting the encoding and segmentation of continuous experience into discrete events, making them particularly relevant for investigating oscillatory dynamics underlying event memory.

### Intracranial EEG preprocessing

To facilitate comparisons of oscillatory activity across brain regions, data was re-referenced using a common average reference scheme, which reduces spatially widespread noise shared across electrodes, and sampled to 512Hz. An automated event-level artifact was applied to remove system-level line noise, eye-blink artifacts, sharp transients, and interictal epileptiform discharges (IEDs). To do so, we calculated a z-score for every time point based on the gradient (first derivative) on the amplitude after applying a 250 Hz high-pass filter. Any time point that exceeded a z-score of 5 with either gradient or high frequency amplitude was marked as artificial, including periods of 200ms before and after each identified time point. Time periods classified as pathological or contaminated by noise were marked as NaN and omitted from further analyses. Because the analysis targeted neural activity around event boundaries, we excluded channels in which the identified events occurred in over 20% of event boundaries, to avoid contamination of boundary-related estimates.

### Spectral Power Analysis

Time-resolved oscillatory power was estimated using a continuous wavelet transform (CWT) with complex Morlet wavelets (wave number = 6). Analyses were performed using frequencies logarithmically spaced between 1 and 200 Hz (192 frequency steps). Power was computed as the squared magnitude of the wavelet coefficients and expressed in decibels, followed by normalization by subtracting the mean power across time within each frequency. To capture finer temporal dynamics, event duration was represented at a second-by-second resolution, where power was computed by averaging within bins of 1s after excluding samples contaminated by interictal epileptiform discharges (IEDs). Bins were retained only if at least 50% of samples were free of IEDs; otherwise, values were set to missing. Power dynamics within each event were quantified by fitting a linear regression to the time-resolved power values, with the resulting slope used as an index of within-event change. In addition, peri-boundary activity was averaged using ±1 s windows around event onset and offset. This was motivated by the fact that, in the current movie stimuli, event boundaries are shaped by cinematic editing conventions and are therefore often only approximately defined in time. For example, auditory cues from an upcoming scene may precede the visual transition (e.g., audio bridging across cuts), which can introduce uncertainty regarding the precise moment at which participants perceptually experience an event boundary. For each frequency band, power values were averaged across frequencies to obtain band-limited power estimates.

### Phase Synchrony Analysis

Phase-based functional connectivity between hippocampal and cortical regions was quantified using the weighted phase-lag index (wPLI; Vinck *et al.*, 2011) in sliding time windows, as follows:

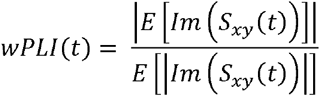

where *S_xy_* is the complex cross-spectrum between signals *x* and *y* and *lm*(*S_xy_*) its imaginary part. The weighted equation was used as it minimizes the influence of volume conduction and common reference effects by selectively weighting consistent non-zero phase lags while down-weighting interactions with phase differences close to zero. wPLI values range from 0 to 1, with higher values reflecting stronger consistency of non-zero phase-lagged synchronization across observations, and values near 0 indicating either weak or inconsistent phase relationships.

Broadband signals were decomposed using Morlet wavelet transforms across logarithmically spaced frequencies between 1 and 200 Hz (192 frequency steps), and instantaneous phase was extracted via the analytic signal (Hilbert transform). The number of wavelet cycles was adjusted as a function of frequency to maintain a constant temporal resolution (FWHM = 0.2 s). This choice is relevant for synchrony analyses because phase-based connectivity measures such as wPLI are sensitive to temporal smoothing; keeping the effective window size constant across frequencies ensures that differences in synchrony are not driven by frequency-dependent changes in temporal averaging. Connectivity was computed between all hippocampal and cortical channel pairs within 2s windows, advanced in 1s steps, yielding a time-resolved estimate of phase synchronization while minimizing zero-lag coupling effects (Fig 1d). Windows containing interictal epileptiform discharges (IEDs) in either region were set to NaN to avoid contamination by pathological activity. Similar to the power analysis, connectivity was quantified in relation to event structure. Each event was segmented into a start period (±1 s around event onset), an end period (±1 s around event offset), and a within event period spanning the event duration where a line was fitted using a linear regression so that the resulting slope could be used as an index of within-event change. For each frequency band, wPLI values were averaged across frequencies to obtain band-limited connectivity estimates.

### Phase Reset Analysis

Event boundary phase resets were quantified using a time–frequency decomposition based on complex Morlet wavelets. Oscillatory activity was estimated across logarithmically spaced frequencies between 1 and 200 Hz (192 frequency steps, wave number = 6), and instantaneous phase was extracted from the analytic signal. Phase estimates were extracted within a ±1 s window around each event boundary annotation. To ensure that phase estimates were not contaminated by pathological activity, any boundary window containing interictal epileptiform discharges (IEDs) was set to NaN. Phase consistency was then quantified using inter-trial phase coherence (ITPC; Makeig *et al.*, 2004), computed as the magnitude of the average unit-phase vector across boundaries, as follows:

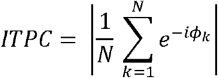

Where N corresponds to the number of trials and *Φ_t_* to the phase at trial *k*. A debiased estimator was additionally used to reduce bias due to finite sample size (84), especially relevant when dividing the trials between recalled and forgotten events:

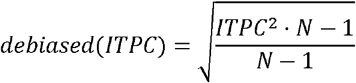

Debiased ITPC values are bounded between 0 and 1, where values close to 0 indicate weak or inconsistent phase alignment across trials, and values close to 1 reflect strong phase consistency (i.e., reliable phase locking across trials). ITPC was computed for all events, and separately for subsets of subsequently recalled and forgotten events, allowing comparison of phase alignment as a function of memory outcome. In addition, null distributions were generated using permutation-based shuffling of event timing to assess whether observed phase consistency exceeded chance levels under temporally randomized boundary assignments (Fig 1e).

### Linear Mixed Models

To assess the relationship between oscillatory activity and memory performance, linear mixed-effects models (LMMs) were fitted separately for each frequency band and brain region. All analyses were performed at the trial level, with observations aggregated across channels within each region. Models were implemented using a random-intercept structure with participant as the grouping factor to account for within-subject dependencies arising from repeated measures across trials and electrodes. For the phase synchrony analyses, the models additionally included crossed random effects for individual electrodes contributing to each pairwise connection. Specifically, variance components were specified for both channels forming each connection (subject × channel identity), in addition to a random intercept for participants, to account for non-independence arising from repeated sampling of electrodes across multiple pairs. Trial duration was included as a fixed-effect covariate in all models to account for variability in event length across trials.

First, band-specific estimates were entered into LMMs predicting power or synchrony as a function of subsequent memory outcome. Specifically, we tested the effects of memory for the subsequent event and the preceding event, as well as their interaction (*memory_stbseq_* * *memory_Qrev_*). Memory variables were coded as binary categorical predictors (recalled vs. forgotten). This model allowed us to assess whether oscillatory activity was related to encoding success for upcoming and past events, and whether these effects interacted.

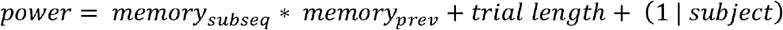

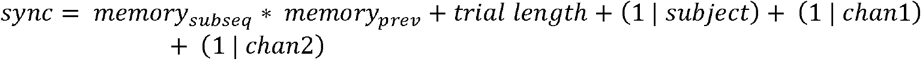

Second, to characterize within-event temporal dynamics, slope estimates were entered into separate LMMs predicting slope as a function of subsequent memory (recalled vs. forgotten).

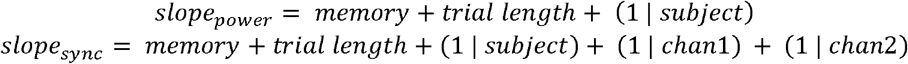

This approach tested whether systematic increases or decreases in oscillatory power or synchrony over the course of an event were associated with later memory performance.

Third, to test whether the boundary-related memory effects differed between the event that had just ended and the event about to begin, we used the same model, which included memory for the preceding event, memory for the following event, and their interaction, and tested a linear contrast between the two memory effects. Specifically, we used a t-test on the fixed-effect estimates to test whether the coefficient for memory of the preceding event minus the coefficient for memory of the following event differed from zero. A significant contrast indicates that boundary activity predicts memory for the preceding and the following event differently, whereas a non-significant contrast indicates no evidence for such a difference:

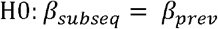

In all models, statistical inference was based on maximum likelihood estimation, and effects were evaluated separately for each frequency band and anatomical region. To enable comparisons across frequency bands and regions, p-values were Bonferroni-corrected for the six frequency bands and three regions tested within each analysis.

## Acknowledgments

This work was supported by the Spanish Ministerio de Ciencia, Innovación y Universidades, which is part of Agencia Estatal de Investigación (AEI), through the project PID2019-111199GB-I00 and PID2022-140426NB-I00 to L.F. (Co-funded by European Regional Development Fund. ERDF, a way to build Europe). We thank the CERCA Programme/Generalitat de Catalunya for institutional support. The project that gave rise to these results received the support of a fellowship of M.Si. from the Human Frontier Science Program (HFSP). The fellowship code is “LT0037/2024-L”.

## Author Contributions Statement

M.Si. and L.F. designed the research; M.Si, X.W. and M.Sa collected the data; E.C, P.R., A.D. and M.C implanted electrodes; M.Si., C.B and J.J. analyzed data; M.Si., C.B., J.J. discussed the results; M.Si., C.B. and J.J wrote the paper. All authors reviewed and revised the final manuscript.

