## Supplementary Material for "Dynamic hippocampal and cortical signatures of naturalistic event memory"

**Table S1:** **Linear mixed-effects model results for the relationship between power and memory performance at boundaries.** Linear mixed-effects models were fitted separately for each frequency band and brain region, with power as the dependent variable and subsequent-event memory (Beginning), preceding-event memory (End), their interaction (Beginning × End), and trial duration as fixed effects. The table reports the estimated regression coefficient (Coef), standard error (SE), uncorrected p-value, Bonferroni-corrected p-value, and significance status for each predictor. Models were considered converged when the fitting procedure successfully reached convergence. The Beg Vs End predictor is the contrast between the Beginning and End coefficients.

| **Region** | **Band** | **Predictor** | **Coef** | **SE** | **pval** | **Converged** | **pval_bon** | **Sig** |
| --- | --- | --- | --- | --- | --- | --- | --- | --- |
| **hipp** | Slow Theta (2–5 Hz) | Beginning | -0.082 | 0.136 | 0.548 | TRUE | 1.000 |  |
| **hipp** | Slow Theta (2–5 Hz) | End | 0.543 | 0.139 | 0.000 | TRUE | 0.002 | ** |
| **hipp** | Slow Theta (2–5 Hz) | Interaction | -0.432 | 0.216 | 0.046 | TRUE | 0.820 |  |
| **hipp** | Slow Theta (2–5 Hz) | Beg Vs End | 0.624 | 0.164 | 0.000 | TRUE | 0.003 | ** |
| **hipp** | Fast Theta (5–10 Hz) | Beginning | 0.226 | 0.115 | 0.050 | TRUE | 0.894 |  |
| **hipp** | Fast Theta (5–10 Hz) | End | 0.384 | 0.118 | 0.001 | TRUE | 0.020 | * |
| **hipp** | Fast Theta (5–10 Hz) | Interaction | -0.338 | 0.183 | 0.065 | TRUE | 1.000 |  |
| **hipp** | Fast Theta (5–10 Hz) | Beg Vs End | 0.157 | 0.139 | 0.258 | TRUE | 1.000 |  |
| **hipp** | Alpha (10–15 Hz) | Beginning | 0.410 | 0.095 | 0.000 | TRUE | 0.000 | *** |
| **hipp** | Alpha (10–15 Hz) | End | 0.184 | 0.097 | 0.056 | TRUE | 1.000 |  |
| **hipp** | Alpha (10–15 Hz) | Interaction | -0.499 | 0.151 | 0.001 | TRUE | 0.017 | * |
| **hipp** | Alpha (10–15 Hz) | Beg Vs End | -0.225 | 0.115 | 0.049 | TRUE | 0.883 |  |
| **hipp** | Beta (15–30 Hz) | Beginning | 0.120 | 0.063 | 0.058 | TRUE | 1.000 |  |
| **hipp** | Beta (15–30 Hz) | End | -0.104 | 0.065 | 0.106 | TRUE | 1.000 |  |
| **hipp** | Beta (15–30 Hz) | Interaction | -0.063 | 0.101 | 0.533 | TRUE | 1.000 |  |
| **hipp** | Beta (15–30 Hz) | Beg Vs End | -0.224 | 0.076 | 0.003 | TRUE | 0.061 |  |
| **hipp** | Slow Gamma (30–55 Hz) | Beginning | -0.029 | 0.046 | 0.532 | TRUE | 1.000 |  |
| **hipp** | Slow Gamma (30–55 Hz) | End | -0.011 | 0.047 | 0.815 | TRUE | 1.000 |  |
| **hipp** | Slow Gamma (30–55 Hz) | Interaction | 0.009 | 0.073 | 0.898 | TRUE | 1.000 |  |
| **hipp** | Slow Gamma (30–55 Hz) | Beg Vs End | 0.018 | 0.055 | 0.749 | TRUE | 1.000 |  |
| **hipp** | Fast Gamma (>60 Hz) | Beginning | -0.162 | 0.034 | 0.000 | TRUE | 0.000 | *** |
| **hipp** | Fast Gamma (>60 Hz) | End | -0.138 | 0.035 | 0.000 | TRUE | 0.001 | ** |
| **hipp** | Fast Gamma (>60 Hz) | Interaction | 0.209 | 0.055 | 0.000 | TRUE | 0.002 | ** |
| **hipp** | Fast Gamma (>60 Hz) | Beg Vs End | 0.023 | 0.041 | 0.572 | TRUE | 1.000 |  |
| **temp** | Slow Theta (2–5 Hz) | Beginning | 0.234 | 0.058 | 0.000 | TRUE | 0.001 | *** |
| **temp** | Slow Theta (2–5 Hz) | End | -0.278 | 0.061 | 0.000 | TRUE | 0.000 | *** |
| **temp** | Slow Theta (2–5 Hz) | Interaction | 0.289 | 0.093 | 0.002 | TRUE | 0.034 | * |
| **temp** | Slow Theta (2–5 Hz) | Beg Vs End | -0.513 | 0.068 | 0.000 | TRUE | 0.000 | *** |
| **temp** | Fast Theta (5–10 Hz) | Beginning | 0.070 | 0.051 | 0.172 | TRUE | 1.000 |  |
| **temp** | Fast Theta (5–10 Hz) | End | -0.358 | 0.053 | 0.000 | TRUE | 0.000 | *** |
| **temp** | Fast Theta (5–10 Hz) | Interaction | 0.494 | 0.081 | 0.000 | TRUE | 0.000 | *** |
| **temp** | Fast Theta (5–10 Hz) | Beg Vs End | -0.427 | 0.060 | 0.000 | TRUE | 0.000 | *** |
| **temp** | Alpha (10–15 Hz) | Beginning | 0.104 | 0.042 | 0.013 | TRUE | 0.240 |  |
| **temp** | Alpha (10–15 Hz) | End | -0.188 | 0.044 | 0.000 | TRUE | 0.000 | *** |
| **temp** | Alpha (10–15 Hz) | Interaction | 0.272 | 0.067 | 0.000 | TRUE | 0.001 | *** |
| **temp** | Alpha (10–15 Hz) | Beg Vs End | -0.293 | 0.050 | 0.000 | TRUE | 0.000 | *** |
| **temp** | Beta (15–30 Hz) | Beginning | -0.071 | 0.027 | 0.009 | TRUE | 0.158 |  |
| **temp** | Beta (15–30 Hz) | End | -0.013 | 0.028 | 0.636 | TRUE | 1.000 |  |
| **temp** | Beta (15–30 Hz) | Interaction | 0.148 | 0.043 | 0.001 | TRUE | 0.011 | * |
| **temp** | Beta (15–30 Hz) | Beg Vs End | 0.057 | 0.032 | 0.071 | TRUE | 1.000 |  |
| **temp** | Slow Gamma (30–55 Hz) | Beginning | -0.250 | 0.023 | 0.000 | FALSE | 0.000 | *** |
| **temp** | Slow Gamma (30–55 Hz) | End | 0.099 | 0.024 | 0.000 | FALSE | 0.000 | *** |
| **temp** | Slow Gamma (30–55 Hz) | Interaction | 0.100 | 0.036 | 0.005 | FALSE | 0.098 |  |
| **temp** | Slow Gamma (30–55 Hz) | Beg Vs End | 0.349 | 0.027 | 0.000 | FALSE | 0.000 | *** |
| **temp** | Fast Gamma (>60 Hz) | Beginning | -0.253 | 0.030 | 0.000 | TRUE | 0.000 | *** |
| **temp** | Fast Gamma (>60 Hz) | End | 0.016 | 0.032 | 0.619 | TRUE | 1.000 |  |
| **temp** | Fast Gamma (>60 Hz) | Interaction | 0.327 | 0.048 | 0.000 | TRUE | 0.000 | *** |
| **temp** | Fast Gamma (>60 Hz) | Beg Vs End | 0.269 | 0.036 | 0.000 | TRUE | 0.000 | *** |
| **front** | Slow Theta (2–5 Hz) | Beginning | 0.023 | 0.059 | 0.692 | TRUE | 1.000 |  |
| **front** | Slow Theta (2–5 Hz) | End | 0.217 | 0.063 | 0.001 | TRUE | 0.011 | * |
| **front** | Slow Theta (2–5 Hz) | Interaction | 0.295 | 0.101 | 0.003 | TRUE | 0.061 |  |
| **front** | Slow Theta (2–5 Hz) | Beg Vs End | 0.193 | 0.071 | 0.007 | TRUE | 0.122 |  |
| **front** | Fast Theta (5–10 Hz) | Beginning | -0.178 | 0.046 | 0.000 | TRUE | 0.002 | ** |
| **front** | Fast Theta (5–10 Hz) | End | -0.109 | 0.049 | 0.026 | TRUE | 0.472 |  |
| **front** | Fast Theta (5–10 Hz) | Interaction | 0.167 | 0.078 | 0.032 | TRUE | 0.572 |  |
| **front** | Fast Theta (5–10 Hz) | Beg Vs End | 0.070 | 0.055 | 0.207 | TRUE | 1.000 |  |
| **front** | Alpha (10–15 Hz) | Beginning | -0.279 | 0.045 | 0.000 | TRUE | 0.000 | *** |
| **front** | Alpha (10–15 Hz) | End | -0.095 | 0.048 | 0.046 | TRUE | 0.832 |  |
| **front** | Alpha (10–15 Hz) | Interaction | 0.298 | 0.076 | 0.000 | TRUE | 0.002 | ** |
| **front** | Alpha (10–15 Hz) | Beg Vs End | 0.184 | 0.054 | 0.001 | TRUE | 0.011 | * |
| **front** | Beta (15–30 Hz) | Beginning | -0.091 | 0.028 | 0.001 | TRUE | 0.021 | * |
| **front** | Beta (15–30 Hz) | End | -0.158 | 0.030 | 0.000 | TRUE | 0.000 | *** |
| **front** | Beta (15–30 Hz) | Interaction | 0.153 | 0.048 | 0.002 | TRUE | 0.027 | * |
| **front** | Beta (15–30 Hz) | Beg Vs End | -0.067 | 0.034 | 0.048 | TRUE | 0.869 |  |
| **front** | Slow Gamma (30–55 Hz) | Beginning | -0.080 | 0.027 | 0.003 | TRUE | 0.050 | * |
| **front** | Slow Gamma (30–55 Hz) | End | -0.094 | 0.029 | 0.001 | TRUE | 0.018 | * |
| **front** | Slow Gamma (30–55 Hz) | Interaction | -0.006 | 0.046 | 0.893 | TRUE | 1.000 |  |
| **front** | Slow Gamma (30–55 Hz) | Beg Vs End | -0.014 | 0.032 | 0.665 | TRUE | 1.000 |  |
| **front** | Fast Gamma (>60 Hz) | Beginning | -0.177 | 0.024 | 0.000 | TRUE | 0.000 | *** |
| **front** | Fast Gamma (>60 Hz) | End | -0.058 | 0.026 | 0.027 | TRUE | 0.484 |  |
| **front** | Fast Gamma (>60 Hz) | Interaction | 0.136 | 0.042 | 0.001 | TRUE | 0.019 | * |
| **front** | Fast Gamma (>60 Hz) | Beg Vs End | 0.120 | 0.030 | 0.000 | TRUE | 0.001 | *** |

**Table S2: Linear mixed-effects model results for the association between within-trial power slopes and memory performance. Models were fitted separately for each frequency band and brain region. with within-trial power slope as the dependent variable and memory outcome and trial duration as fixed effects. Memory was coded as a binary predictor (recalled vs. forgotten).** The table reports the estimated regression coefficient (Coef), standard error (SE), uncorrected p-value, Bonferroni-corrected p-value, and significance status for each predictor. Models were considered converged when the fitting procedure successfully reached convergence.

| **Region** | **Band** | **Predictor** | **Coef** | **SE** | **pval** | **Converged** | **pval_bon** | **Sig** |
| --- | --- | --- | --- | --- | --- | --- | --- | --- |
| **hipp** | Slow Theta (2–5 Hz) | memory | -0.335 | 0.126 | 0.008 | TRUE | 0.048 | * |
| **hipp** | Fast Theta (5–10 Hz) | memory | -0.164 | 0.116 | 0.156 | TRUE | 0.936 |  |
| **hipp** | Alpha (10–15 Hz) | memory | -0.122 | 0.099 | 0.217 | TRUE | 1.000 |  |
| **hipp** | Beta (15–30 Hz) | memory | -0.194 | 0.057 | 0.001 | TRUE | 0.004 | ** |
| **hipp** | Slow Gamma (30–55 Hz) | memory | 0.033 | 0.040 | 0.402 | TRUE | 1.000 |  |
| **hipp** | Fast Gamma (>60 Hz) | memory | 0.024 | 0.031 | 0.443 | TRUE | 1.000 |  |
| **temp** | Slow Theta (2–5 Hz) | memory | -0.629 | 0.051 | 0.000 | TRUE | 0.000 | *** |
| **temp** | Fast Theta (5–10 Hz) | memory | -0.446 | 0.044 | 0.000 | TRUE | 0.000 | *** |
| **temp** | Alpha (10–15 Hz) | memory | -0.336 | 0.037 | 0.000 | TRUE | 0.000 | *** |
| **temp** | Beta (15–30 Hz) | memory | -0.101 | 0.024 | 0.000 | TRUE | 0.000 | *** |
| **temp** | Slow Gamma (30–55 Hz) | memory | 0.206 | 0.020 | 0.000 | TRUE | 0.000 | *** |
| **temp** | Fast Gamma (>60 Hz) | memory | 0.249 | 0.029 | 0.000 | TRUE | 0.000 | *** |
| **front** | Slow Theta (2–5 Hz) | memory | -0.261 | 0.057 | 0.000 | TRUE | 0.000 | *** |
| **front** | Fast Theta (5–10 Hz) | memory | 0.026 | 0.045 | 0.568 | TRUE | 1.000 |  |
| **front** | Alpha (10–15 Hz) | memory | 0.227 | 0.044 | 0.000 | TRUE | 0.000 | *** |
| **front** | Beta (15–30 Hz) | memory | -0.094 | 0.026 | 0.000 | FALSE | 0.002 | ** |
| **front** | Slow Gamma (30–55 Hz) | memory | -0.046 | 0.022 | 0.038 | TRUE | 0.228 |  |
| **front** | Fast Gamma (>60 Hz) | memory | 0.022 | 0.024 | 0.372 | TRUE | 1.000 |  |

**Table S3:** **Linear mixed-effects model results for the relationship between synchrony and memory performance at boundaries.** Linear mixed-effects models were fitted separately for each frequency band and brain region. with synchrony as the dependent variable and subsequent-event memory (Beginning), preceding-event memory (End), their interaction (Beginning × End), and trial duration as fixed effects. The table reports the estimated regression coefficient (Coef), standard error (SE), uncorrected p-value, Bonferroni-corrected p-value, and significance status for each predictor. Models were considered converged when the fitting procedure successfully reached convergence. The Beg Vs End predictor is the contrast between the Beginning and End coefficients.

| **Region** | **Band** | **Predictor** | **Coef** | **SE** | **pval** | **Converged** | **pval_bon** | **Sig** |
| --- | --- | --- | --- | --- | --- | --- | --- | --- |
| **hipp-temp** | Slow Theta (2–5 Hz) | Beginning | -0.005 | 0.003 | 0.064 | TRUE | 0.770 |  |
| **hipp-temp** | Slow Theta (2–5 Hz) | End | -0.014 | 0.003 | 0.000 | TRUE | 0.000 | *** |
| **hipp-temp** | Slow Theta (2–5 Hz) | Interaction | 0.032 | 0.004 | 0.000 | TRUE | 0.000 | *** |
| **hipp-temp** | Slow Theta (2–5 Hz) | Beg Vs End | -0.010 | 0.003 | 0.001 | TRUE | 0.018 | * |
| **hipp-temp** | Fast Theta (5–10 Hz) | Beginning | 0.004 | 0.002 | 0.080 | TRUE | 0.963 |  |
| **hipp-temp** | Fast Theta (5–10 Hz) | End | 0.004 | 0.002 | 0.079 | TRUE | 0.948 |  |
| **hipp-temp** | Fast Theta (5–10 Hz) | Interaction | 0.004 | 0.003 | 0.298 | TRUE | 1.000 |  |
| **hipp-temp** | Fast Theta (5–10 Hz) | Beg Vs End | 0.000 | 0.003 | 0.970 | TRUE | 1.000 |  |
| **hipp-temp** | Alpha (10–15 Hz) | Beginning | 0.005 | 0.002 | 0.013 | TRUE | 0.155 |  |
| **hipp-temp** | Alpha (10–15 Hz) | End | 0.011 | 0.002 | 0.000 | TRUE | 0.000 | *** |
| **hipp-temp** | Alpha (10–15 Hz) | Interaction | -0.001 | 0.003 | 0.846 | TRUE | 1.000 |  |
| **hipp-temp** | Alpha (10–15 Hz) | Beg Vs End | 0.005 | 0.003 | 0.045 | TRUE | 0.545 |  |
| **hipp-temp** | Beta (15–30 Hz) | Beginning | 0.004 | 0.001 | 0.010 | FALSE | 0.120 |  |
| **hipp-temp** | Beta (15–30 Hz) | End | -0.003 | 0.001 | 0.038 | FALSE | 0.459 |  |
| **hipp-temp** | Beta (15–30 Hz) | Interaction | 0.007 | 0.002 | 0.001 | FALSE | 0.008 | ** |
| **hipp-temp** | Beta (15–30 Hz) | Beg Vs End | -0.007 | 0.002 | 0.000 | FALSE | 0.001 | *** |
| **hipp-temp** | Slow Gamma (30–55 Hz) | Beginning | -0.004 | 0.001 | 0.001 | TRUE | 0.016 | * |
| **hipp-temp** | Slow Gamma (30–55 Hz) | End | 0.005 | 0.001 | 0.000 | TRUE | 0.001 | ** |
| **hipp-temp** | Slow Gamma (30–55 Hz) | Interaction | 0.016 | 0.002 | 0.000 | TRUE | 0.000 | *** |
| **hipp-temp** | Slow Gamma (30–55 Hz) | Beg Vs End | 0.008 | 0.001 | 0.000 | TRUE | 0.000 | *** |
| **hipp-temp** | Fast Gamma (>60 Hz) | Beginning | -0.004 | 0.001 | 0.000 | FALSE | 0.000 | *** |
| **hipp-temp** | Fast Gamma (>60 Hz) | End | -0.001 | 0.001 | 0.019 | FALSE | 0.228 |  |
| **hipp-temp** | Fast Gamma (>60 Hz) | Interaction | 0.020 | 0.001 | 0.000 | FALSE | 0.000 | *** |
| **hipp-temp** | Fast Gamma (>60 Hz) | Beg Vs End | 0.002 | 0.001 | 0.002 | FALSE | 0.024 | * |
| **hipp-front** | Slow Theta (2–5 Hz) | Beginning | 0.009 | 0.003 | 0.003 | TRUE | 0.033 | * |
| **hipp-front** | Slow Theta (2–5 Hz) | End | -0.002 | 0.003 | 0.620 | TRUE | 1.000 |  |
| **hipp-front** | Slow Theta (2–5 Hz) | Interaction | 0.007 | 0.005 | 0.138 | TRUE | 1.000 |  |
| **hipp-front** | Slow Theta (2–5 Hz) | Beg Vs End | -0.010 | 0.004 | 0.006 | TRUE | 0.070 |  |
| **hipp-front** | Fast Theta (5–10 Hz) | Beginning | 0.006 | 0.003 | 0.022 | TRUE | 0.267 |  |
| **hipp-front** | Fast Theta (5–10 Hz) | End | 0.007 | 0.003 | 0.018 | TRUE | 0.211 |  |
| **hipp-front** | Fast Theta (5–10 Hz) | Interaction | -0.011 | 0.004 | 0.012 | TRUE | 0.143 |  |
| **hipp-front** | Fast Theta (5–10 Hz) | Beg Vs End | 0.001 | 0.003 | 0.817 | TRUE | 1.000 |  |
| **hipp-front** | Alpha (10–15 Hz) | Beginning | 0.009 | 0.003 | 0.000 | TRUE | 0.003 | ** |
| **hipp-front** | Alpha (10–15 Hz) | End | 0.006 | 0.003 | 0.029 | TRUE | 0.351 |  |
| **hipp-front** | Alpha (10–15 Hz) | Interaction | -0.019 | 0.004 | 0.000 | TRUE | 0.000 | *** |
| **hipp-front** | Alpha (10–15 Hz) | Beg Vs End | -0.003 | 0.003 | 0.309 | TRUE | 1.000 |  |
| **hipp-front** | Beta (15–30 Hz) | Beginning | 0.006 | 0.002 | 0.000 | TRUE | 0.002 | ** |
| **hipp-front** | Beta (15–30 Hz) | End | -0.001 | 0.002 | 0.773 | TRUE | 1.000 |  |
| **hipp-front** | Beta (15–30 Hz) | Interaction | -0.012 | 0.003 | 0.000 | TRUE | 0.000 | *** |
| **hipp-front** | Beta (15–30 Hz) | Beg Vs End | -0.007 | 0.002 | 0.001 | TRUE | 0.015 | * |
| **hipp-front** | Slow Gamma (30–55 Hz) | Beginning | -0.004 | 0.001 | 0.000 | TRUE | 0.006 | ** |
| **hipp-front** | Slow Gamma (30–55 Hz) | End | -0.005 | 0.001 | 0.000 | TRUE | 0.002 | ** |
| **hipp-front** | Slow Gamma (30–55 Hz) | Interaction | 0.006 | 0.002 | 0.006 | TRUE | 0.069 |  |
| **hipp-front** | Slow Gamma (30–55 Hz) | Beg Vs End | -0.001 | 0.002 | 0.638 | TRUE | 1.000 |  |
| **hipp-front** | Fast Gamma (>60 Hz) | Beginning | 0.003 | 0.001 | 0.000 | TRUE | 0.001 | *** |
| **hipp-front** | Fast Gamma (>60 Hz) | End | -0.001 | 0.001 | 0.041 | TRUE | 0.489 |  |
| **hipp-front** | Fast Gamma (>60 Hz) | Interaction | 0.000 | 0.001 | 0.717 | TRUE | 1.000 |  |
| **hipp-front** | Fast Gamma (>60 Hz) | Beg Vs End | -0.004 | 0.001 | 0.000 | TRUE | 0.000 | *** |

**Table S4**: **Linear mixed-effects model results for the association between within-trial synchrony slopes and memory performance. Models were fitted separately for each frequency band and brain region. with within-trial power slope as the dependent variable and memory outcome and trial duration as fixed effects. Memory was coded as a binary predictor (recalled vs. forgotten).** The table reports the estimated regression coefficient (Coef), standard error (SE), uncorrected p-value, Bonferroni-corrected p-value, and significance status for each predictor. Models were considered converged when the fitting procedure successfully reached convergence.

| **Region** | **Band** | **Predictor** | **Coef** | **SE** | **pval** | **Converged** | **pval_bon** | **Sig** |
| --- | --- | --- | --- | --- | --- | --- | --- | --- |
| **hipp-temp** | Slow Theta (2–5 Hz) | memory | -0.001 | 0.002 | 0.396 | TRUE | 1.000 |  |
| **hipp-temp** | Fast Theta (5–10 Hz) | memory | -0.004 | 0.001 | 0.012 | TRUE | 0.070 |  |
| **hipp-temp** | Alpha (10–15 Hz) | memory | -0.004 | 0.001 | 0.002 | TRUE | 0.011 | ** |
| **hipp-temp** | Beta (15–30 Hz) | memory | 0.001 | 0.001 | 0.200 | TRUE | 1.000 |  |
| **hipp-temp** | Slow Gamma (30–55 Hz) | memory | 0.001 | 0.001 | 0.203 | FALSE | 1.000 |  |
| **hipp-temp** | Fast Gamma (>60 Hz) | memory | 0.002 | 0.000 | 0.000 | TRUE | 0.000 | *** |
| **hipp-front** | Slow Theta (2–5 Hz) | memory | -0.008 | 0.002 | 0.000 | TRUE | 0.001 | *** |
| **hipp-front** | Fast Theta (5–10 Hz) | memory | -0.004 | 0.002 | 0.014 | FALSE | 0.086 |  |
| **hipp-front** | Alpha (10–15 Hz) | memory | -0.001 | 0.002 | 0.615 | TRUE | 1.000 |  |
| **hipp-front** | Beta (15–30 Hz) | memory | -0.003 | 0.001 | 0.015 | TRUE | 0.090 |  |
| **hipp-front** | Slow Gamma (30–55 Hz) | memory | -0.002 | 0.001 | 0.011 | FALSE | 0.068 |  |
| **hipp-front** | Fast Gamma (>60 Hz) | memory | 0.001 | 0.000 | 0.021 | TRUE | 0.126 |  |

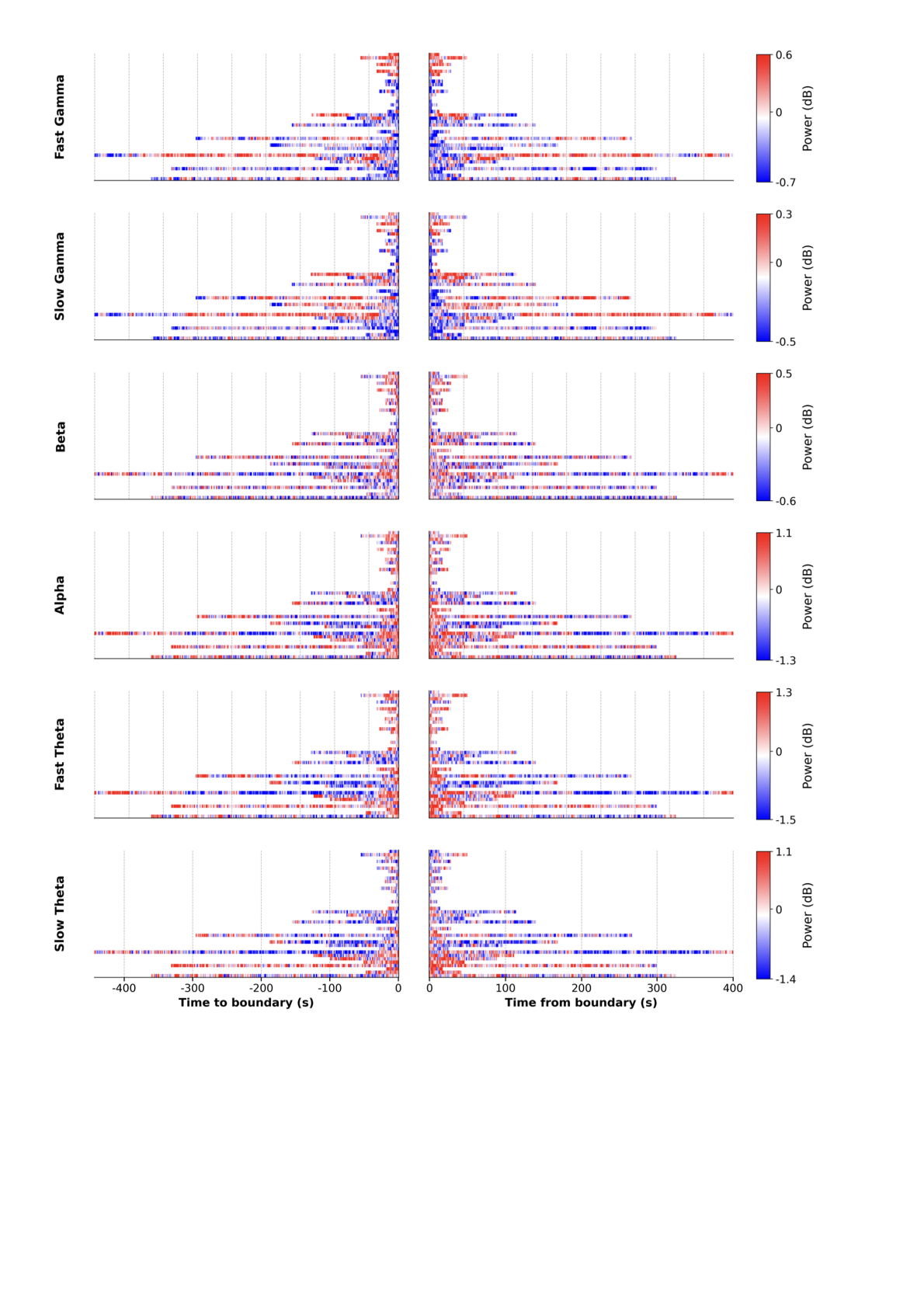

**Figure S1:** **Single-trial power dynamics in an example region (temporal cortex).** Power was averaged within consecutive 1-s bins and is shown for each time bin (x-axis) and each trial (y-axis), separately for every frequency band. Trials are aligned to the end of an event (left column) or to the beginning of an event (right column). The figure illustrates the dynamic structure of the data, including variability in trial length and band-specific power dynamics that differ across positions within an event.

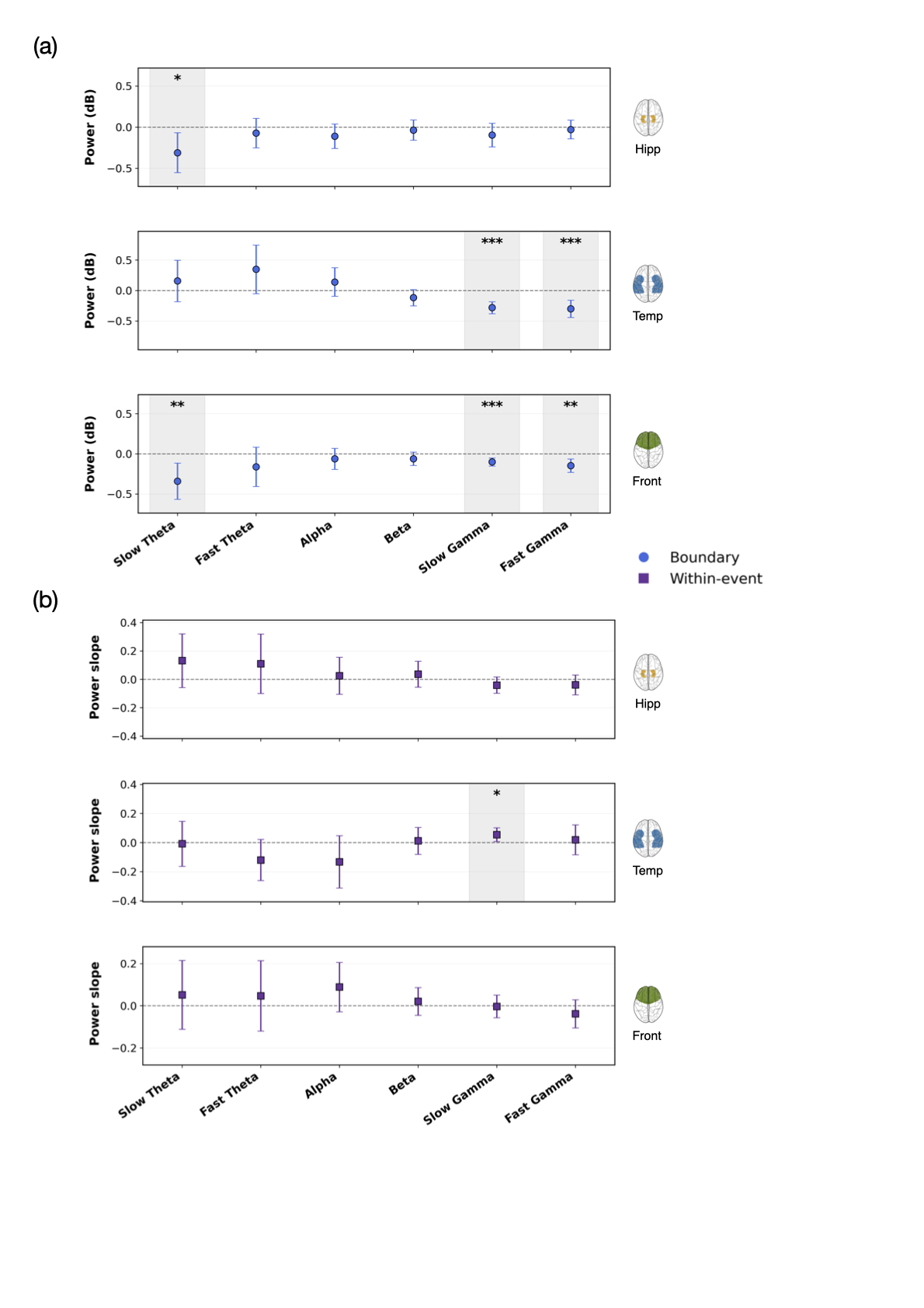

**Figure S2: Overall power modulation at event boundaries and within-event power slopes.** (a) Mean power at boundaries and (b) mean within-event power slope, shown separately for each frequency band (x-axis) and brain region. Slopes were obtained by fitting a linear regression to the time-resolved power values within each event. Error bars show 95% confidence intervals computed from participant-level data. Significance was assessed by testing the values against zero with a one-sample t-test: one, two or three asterisks denote p < 0.05, p < 0.01 and p < 0.001, respectively (Bonferroni-corrected across frequency bands and regions). Shaded grey areas highlight frequency bands with a significant effect.
